# Systematic identification of tissue-conserved m^6^A sites reveals a stable epitranscriptomic regulatory layer controlling essential genes

**DOI:** 10.64898/2026.03.19.713046

**Authors:** Sumin Jo, Tinghe Zhang, Shou-jiang Gao, Yufei Huang

## Abstract

Chemical modifications to RNA are fundamental regulators of cellular identity and function. Among these, N6-methyladenosine (m^6^A) is the most abundant mRNA modification in mammalian cell, governing major post-transcriptional processes. While conditional m^6^A dynamics are well studied, the extent and function of condition-independent, tissue-conserved (TC) m^6^A in humans remain unclear. Here we show that 5,945 TC sites are consistently methylated across 24 human tissues. These sites are enriched near stop codons, evolutionarily conserved, and characterized by distinct sequence signatures. *RBM15/B* are identified as candidate mediators of TC m^6^A deposition, and *YTHDF1-3* and *UPF1* are preferentially enriched at TC sites, supporting their role for m^6^A-linked mRNA decay. TC m^6^A sites mark 1,386 genes essential for core cellular processes like autophagy and homeostasis, showing stable expression and evolutionary constraint. Pan-cancer analysis reveals that TC m^6^A genes are disproportionately differentially expressed, alongside with altered *RBM15/B* expression, suggesting that disruption of this stable m^6^A layer may contribute to transcriptional changes in cancer.

## Introduction

N^6^-methyladenosine (m^6^A) is the most abundant internal modification in eukaryotic mRNA and serves as a fundamental post-transcriptional regulator of gene expression. By governing the life cycle of transcripts - from splicing and nuclear export to translation and decay - this epitranscriptomic layer enables cells to rapidly tune their proteomes in response to developmental cues and environmental stimuli^1^. The m^6^A modification is installed by a writer complex (*METTL3-METTL14*), removed by erasers (*FTO* and *ALKBH5*), and decoded by reader proteins (e.g., *YTHDF1-3*) which mediate various functions such as mRNA degradation, splicing, and translation^2^.

A defining feature of m^6^A is its conditionality: signaling cascades and stress responses can rapidly alter the methylation status of thousands of transcripts, reprogramming pathways involved in development, immunity, and tissue homeostasis^3^. Consequently, the dysregulation of m^6^A machinery is increasingly linked to human diseases, particularly cancer, where aberrant methylation can disrupt the expression of key oncogenes and tumor suppressors^3,4^. Alongside this conditional mode of regulation, emerging evidence suggests the existence of a stable, constitutive layer of m^6^A. Studies in simpler eukaryotes have identified a subset of m^6^A sites that are constitutively methylated, with deposition governed by local sequence features rather than transient cellular states^5,6^, suggesting the existence of a cis-encoded m^6^A program that operates as a fixed property of transcripts. More recently, analysis of m^6^A stoichiometry in mammalian cells showed that highly methylated sites tend to be evolutionarily conserved, providing genetic evidence for a stable m^6^A infrastructure^7^. Existing computational predictions often overestimate this conserved regulation by not accounting for the factors that determine the actual occupancy^8^. More fundamentally, such predictions remain descriptive without elucidating the underlying biological functions. Thus, whether a stable, hard-wired m^6^A layer is broadly conserved across human tissues and if so, what molecular features define it, what biological processes it governs, and how its disruption contributes to disease, remain unknown.

To address these fundamental questions, we performed a systematic, cross-tissue investigation of the human epitranscriptome. By integrating 67 high-quality methylated RNA immunoprecipitation sequencing (MeRIP-seq) datasets spanning 24 normal human tissues, we sought to define a high-confidence set of tissue-conserved (TC) m^6^A sites. Our objective was to move beyond the descriptive catalogs to determine the molecular mechanism that defines these sites, the specific writer and reader architectures that maintain them, and the biological processes they govern. We specifically investigated whether these stable modifications act as a buffered regulatory layer for genes essential to cellular survival and how the disruption of this stability contributes to the transcriptional chaos observed in clinical cancer cohorts. Collectively, our study delineates a consistently methylated subset of the human m^6^A landscape and provides a framework for understanding how the TC m^6^A contributes to conserved post-transcriptional regulation of essential genes, with potential implications for both normal tissue homeostasis and disease.

## Results

### Systematic identification of a stable tissue-conserved human m^6^A landscape

We assembled 67 MeRIP-seq samples spanning 24 human tissue types, comprising 22 samples from five male donors in the Chinese Brain Bank Center (CBBC) dataset^9^ and 45 samples from seven donors (one female and six males) from the National Genomics Data Center (NGDC) dataset^10^ (Fig. 1a). Then applied these MeRIP-samples to a pipeline we developed, which integrates MeRIP-seq data with high-confidence single nucleotide m^6^A sites^11,12^, to identify TC m^6^A sites (Supplementary Fig. 1). m^6^A peaks were called using exomePeak2^13,14^ with stringent filtering to remove low-quality signals (see Methods), yielding 432,343 peaks in the CBBC (mean = 19,652 per sample) and 1,554,830 peaks in the NGDC dataset (mean = 34,552 per sample) (Fig. 1b; Supplementary Fig. 2; Supplementary Table 1). Peak counts differed significantly between datasets (Wilcoxon rank-sum test, *P* = 1.4 × 10⁻¹⁰), likely reflecting variation in sequencing depth and experimental protocol (Supplementary Fig. 3).

**Fig. 1.**
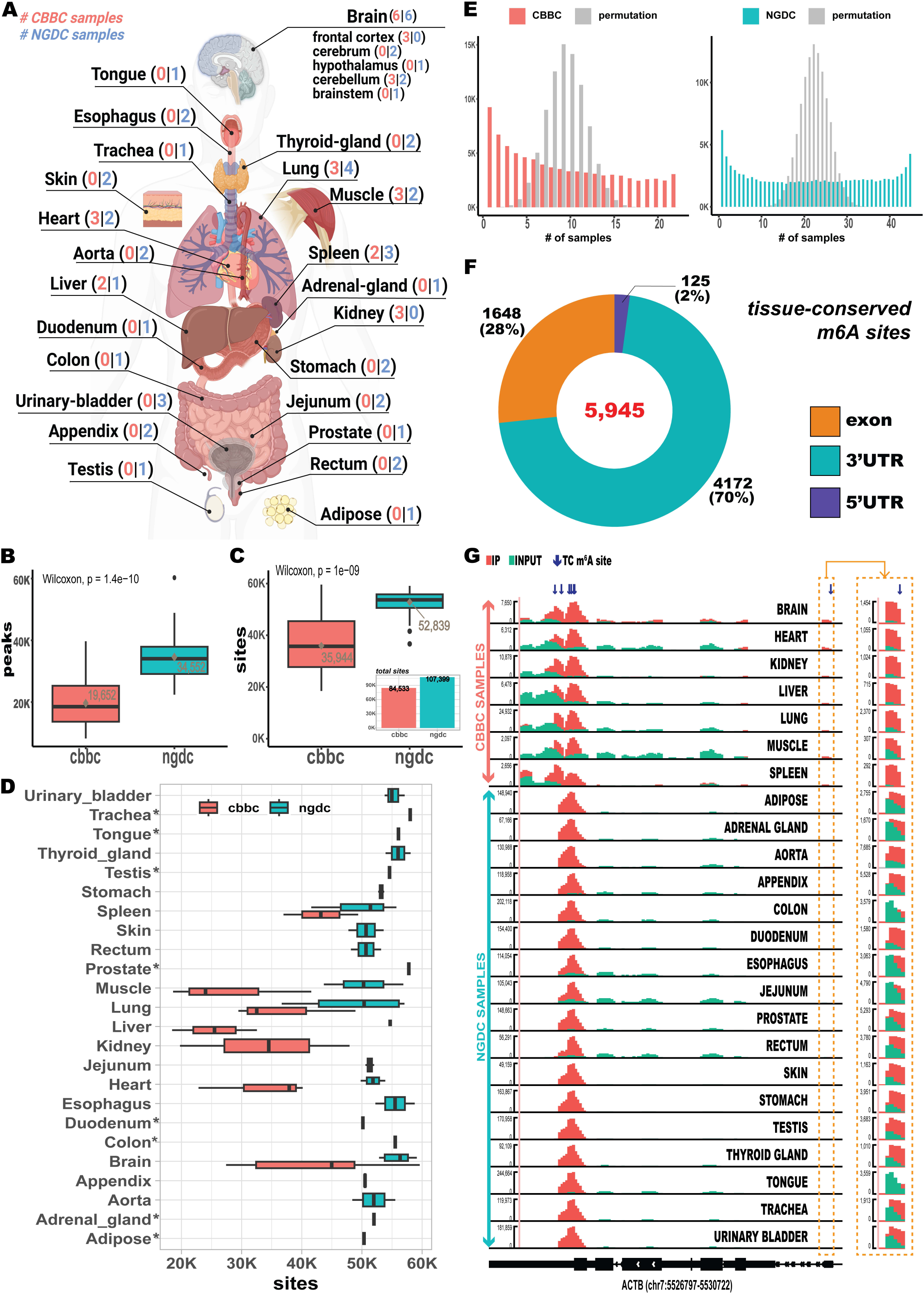
Data overview and identification of TC m6A sites. (A) Anatomical representation of human MeRIP-seq samples used in this study. Number of samples in CBBC and NGDC dataset are indicated by colors respectively. (B) Distribution of predicted m6A peaks across tissue samples (C) Distribution of m6A sites residing within m6A peaks. A total of 84,533 and 107,399 m6A sites were mapped to m6A peaks in the CBBC and NGDC datasets, respectively. (D) Distribution of m6A sites overlapping with m6A peaks in each tissue. Asterisks indicate tissues represented by only a single sample. (E) Distribution of the number of sample overlaps for individual m6A sites compared with the null distribution. (F) Genomic regional distribution of TC m6A sites. (G) IGV tracks of ACTB gene with its TC sites. Green and red tracks indicate transcripts in input control sample and m6A-antibody immunoprecipitated (ip) sample, respectively. The top seven tracks represent CBBC samples; the remaining tracks represent NGDC tissue samples. TC m6A sites are indicated by blue arrows. Track heights are indicated on the right.

To map methylation at nucleotide resolution, we intersected m^6^A peaks with a reference set of 124,291 high-confidence single-base sites, each supported by at least two independent single-base profiling techniques (see Methods). These background sites were predominantly located in 3’ untranslated regions (3’UTRs, 53%), followed by exons (41%) and 5′UTRs (5%), consistent with the canonical m^6^A metagene distribution^15,16^. We then quantified sample-specific methylation events by overlapping called peaks with background sites. This yielded 790,762 m^6^A sites across CBBC samples (mean = 35,944 per sample) and 2,373,750 sites in NGDC (mean = 52,839 per sample) (Fig. 1c, d; Supplementary Table 2). Although the total site number was high, the unique m^6^A sites per dataset were considerably low, (84,533 sites in CBBC and 107,399 sites in NGDC), suggesting the substantial sharing of high-confidence sites across samples. Notably, 82,951 sites (98% of CBBC data) were common to both datasets, pointing to a stable subset of sites. To assess the extent of a site sharing across tissues, we examined the distribution of sample counts per site. Compared to a permutation-based null distribution, we observed a bimodal pattern in both datasets: many sites were unique to one or a few samples, whereas a distinct subset was recurrent across nearly all samples (Fig. 1e). This suggested the coexistence of both tissue-specific and TC methylation programs. Using permutation tests, we identified 13,218 sites in CBBC and 27,692 in NGDC that were significantly shared across samples more than expected by chance (FDR ˂ 0.01). The vast majority (95% in CBBC and 93% in NGDC) were detected in nearly all tissues (Supplementary Fig. 4). In addition, we identified 41,966 sites that were infrequently methylated (i.e., methylated in ≤ 3 tissues) encompassing both CBBC and NGDC tissues. Most of these infrequent sites (82.6%) were only present in fewer than five samples (Supplementary Fig. 5). To derive a robust set of TC m^6^A sites, we intersected the significantly shared m^6^A sites from CBBC and NGDC, and retained only those presented in all 24 tissue types. This resulted in 5,945 high-confidence TC m^6^A sites (FDR ˂ 0.0054) in 1,386 genes (Fig. 1f; Supplementary Table 3). These sites were enriched in 3’UTRs (70%), followed by exons (28%) and 5’UTRs (2%), a distribution similar to that of background sites. They represented 7.2% of the 82,951 sites shared across datasets and 5% of background sites.

To further evaluate their reproducibility and context independence, we analyzed the origin of TC sites across cell lines, profiling techniques, and experimental conditions in the original single-nucleotide datasets (Supplementary Figs. 6 and 7). TC sites were more likely to be detected in ≥ 3 cell lines (58.7% vs. 28.3%), ≥ 3 profiling methods (72.4% vs. 45.8%), ˃ 4 independent studies (50.5% vs. 20.1%), and ˃ 3 perturbation conditions (37.4% vs. 18.9%) than background sites. Despite the overrepresentation of transformed cell lines and perturbed states in original single-base sites data^11,12^, these observations provide additional support for the identified TC sites being tissue conserved. Lastly, we examined their presence in genes known to exhibit stable m^6^A methylation. *ACTB*, a frequently used positive control in m^6^A profiling, harbored seven TC m^6^A sites (Fig. 1g), consistent with its broad and stable methylation across cell conditions ^17,18^. Similarly, we identified four TC m^6^A sites in *BSG*, a standard benchmark for high-confidence m^6^A site validation^19^, and two significantly shared sites in *MALAT1*, where conserved m^6^A residues are established determinants of lncRNA structure and stability^20,21^. Together, these findings define a core set of m^6^A sites that exhibit remarkably stable deposition across diverse human tissues and suggest the presence of a conserved layer of m^6^A regulation operating across human tissues.

### Tissue-conserved m^6^A sites exhibit distinct positional, structural, and evolutionary signatures

We next investigated the molecular characteristics that distinguish TC sites from infrequently methylated and background m^6^A sites. TC sites were significantly more enriched near the stop codon compared to both infrequent and background sites (Fig. 2a) and exhibited a sharper 3’UTR density peak, reinforcing their preferential localization to the canonical m^6^A regulatory region. We then compared the methylation levels and found that TC sites exhibited significantly higher methylation than both infrequent and background sites (Wilcoxon rank-sum test, *P* < 0.001; Fig. 2b), a pattern consistently observed across all 24 tissue types (Supplementary Fig. 8). To determine whether this enrichment could be explained solely by their positional bias, we examined methylation levels across different transcript regions. TC sites maintained significantly higher methylation regardless of genomic location (Supplementary Fig. 9), suggesting that the elevated methylation is not simply a positional artifact but an important feature of TC sites.

**Fig. 2.**
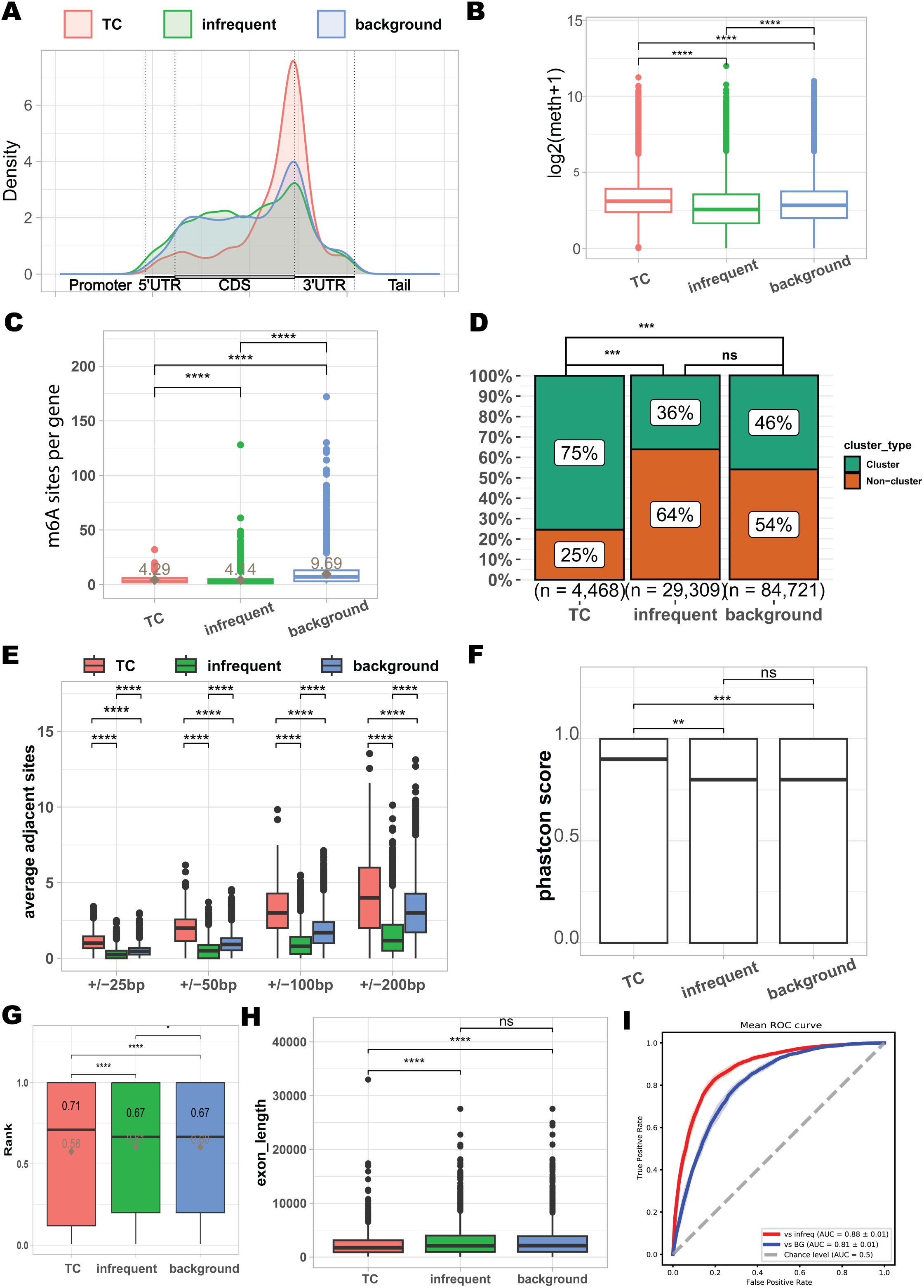
Genomic and architectural features delineate distinct classes of m6A sites. Three categories of m6A sites were defined for analysis: tissue-conserved (TC) sites, infrequent sites, and background sites. (A) meta­gene distribution of TC sites, infrequent sites, and background sites. (B) m6A sites and their peak-methylations where they resided. Square indicates the average and bar in boxplot indicates the median. (C) Distribution of the number of sites per gene for each category. On average, a gene contains 4.29 TC m6A sites, 4.14 infrequent sites, and 9.96 background sites. (D) Enrichment analysis comparing each category to a reference set of clustered m6A sites previously identified by the GLORI method. TC sites show the highest overlap with these high-confidence clusters. (E) Analysis of local site density, showing the average number of adjacent m6A sites within variable window sizes (from ±25 bp to ±200 bp). This measures the degree of local clustering for each category. (F) Distribution of PhastCons conservation scores, demonstrating that TC sites are located in the most evolutionarily conserved regions. (G-H) Analysis of the exon architecture context for each site category. (G) Distribution of a normalized exon rank (rank = exon order / total number of exons) reveals the positional preference of sites within a transcript’s splicing structure. Horizontal bar is median and diamond is mean (H) Distribution of the lengths of exons harboring each category of m6A site. (I) Evaluation of a fine-tuned m6A-BERT sequence model on its ability to classify the intrinsic sites compared with infrequent sites (red line) and background sites (blue line), with performance shown as a Receiver Operating Characteristic (ROC) curve.

Next, we examined the distribution of m^6^A sites per gene. TC sites were present at an average of 4.29 sites per gene, slightly more than infrequent (4.14) but notably fewer than background sites (9.69), indicating a more focused deposition pattern (Fig. 2c). Consistent with a recent study that revealed frequent clustered m^6^A sites^12^, TC sites were significantly enriched among reported clustered sites, compared to both infrequent (Fisher’s exact test, *OR* = 5.40 and *P* < 2.2 × 10⁻¹⁶; Fig. 2d) and background sites (Fisher’s exact test, *OR* = 3.59, *P* < 2.2 × 10⁻¹⁶; Fig. 2d). To investigate whether TC sites exhibit a greater tendency to form clusters, we quantified the number of nearby m^6^A sites within ±25 to ±200 bp windows. TC sites had consistently more adjacent m^6^A sites than infrequent or background sites across all tested distances (Wilcoxon rank-sum test, *P* < 0.0001; Fig. 2e). Within ±200 bp, each TC site was accompanied by an average of 4.25 neighboring TC sites, compared to 1.52 for infrequent and 3.12 for background sites, indicating that the majority of TC sites within a gene are more clustered than the other sites. This spatial proximity may contribute to their stable methylation across tissues.

We next assessed sequence conservation at the nucleotide level, as functional regulatory elements are often evolutionarily constrained^7,9^. All m^6^A sites exhibited positive phyloP scores^22^, consistent with m^6^A localization in conserved or neutrally evolving regions (Supplementary Fig. 10). However, TC sites showed significantly higher phastCons conservation scores^23^ than both infrequent and background sites (Fig. 2f), suggesting that TC sites are more likely to reside in functionally important cis-regulatory contexts.

As m^6^A deposition can be influenced by exon structure and exon junction complexes (EJCs)^24–26^, we tested whether TC sites are less susceptible to EJC-mediated inhibition. We first calculated a normalized positional rank of exon towards transcript end for each site (exon order divided by total number of exons; Fig. 2g). TC sites showed a strong bias toward the 3’ end of transcripts, with a median rank of 0.71. In contrast, infrequent/background sites were located more centrally, with a median rank of 0.67, a difference that was found to be statistically significant (Wilcoxon rank-sum test, *P* < 0.05). Notably, the distribution of TC sites ranks was left-skewed (mean = 0.58), suggesting a bimodal population with a substantial proportion of sites located at the transcript ends (first and last exons). In contrast, the ranks in infrequent/background sites were more symmetrically distributed around the median (mean = 0.6). In addition, we examined the length of the exons harboring these m^6^A sites (Fig. 2h). All three groups were found in exceptionally long exons (mean > 2300 bp). However, compared to infrequent and background sites, TC sites resided in significantly shorter exons on average (mean = 2359bp vs 2895bp (infrequent) and 2813bp (background), respectively). And no significant difference in exon length was observed between infrequent and background sites. Together, TC m^6^A sites preferentially localize to the boundaries of transcripts (primarily the last exon), where the EJC-mediated m^6^A exclusion is weaker than internal region. Infrequent/background sites are situated more centrally within the transcript, in exons that are, on average, significantly longer. To further assess whether TC methylation is associated with cis-regulatory sequence features, we fine-tuned m^6^A-BERT^27^, a model previously optimized for m^6^A site recognition, to classify ±100 bp flanking sequences of TC sites versus those of infrequent or background sites. The model achieved an area under the curve (AUC) of 0.88 and 0.81, respectively (Fig. 2i), indicating that TC sites possess distinct local sequence features that reliably distinguish them from infrequent or background m^6^A sites.

Together, these findings demonstrate that TC m^6^A sites are enriched near stop codons, tend to exhibit higher methylation and greater local clustering, and display elevated conservation relative to infrequent and background sites, with a bias toward terminal exons. Moreover, the ability of a sequence-based model to distinguish TC sites from other site classes suggests that local sequence context may contribute to their reproducible methylation across tissues.

### TC m^6^A deposition is associated with RBM15/B and fine-tuned by upstream RNA-binding proteins

The distinct sequence-context features of TC m^6^A sites raise the possibility that local cis-elements contribute to their stable deposition across tissues. To investigate mechanisms, we first examined whether TC sites show consistent methylation patterns across samples. Across tissue types, TC sites showed similar methylation levels (Fig. 3a), exhibiting significantly higher cosine similarity and lower standard deviation in their methylation levels than expected under a random model (Wilcoxon rank-sum test, *P* < 2.22 × 10⁻¹⁶; Fig. 3b,c; Supplementary Fig. 11). These patterns are consistent with the idea that TC site methylation is shaped by tissue-independent regulatory mechanisms. Canonical m^6^A deposition is catalyzed by the *METTL3-METTL14* methyltransferase complex (MTC), assisted by accessory proteins such as *RBM15, RBM15B, VIRMA, and ZC3H13*^28^. This complex preferentially modifies the DRACH motif (D = A/G/U; R = A/G; H = A/C/U)^28^. To assess whether TC sites conform to this motif, we performed de novo motif analysis using MEME^29,30^. All three site classes including TC, infrequent, and background were enriched for the consensus 5′-RGACW-3′ sequence (R = A/G; W = A/U), a variant of DRACH, with *E*-values of 1.3 × 10⁻² (TC), 6.1 × 10⁻⁴ (infrequent), and 1.6 × 10⁻⁷ (background) (Fig. 3d). Additionally, 92% of TC sites contained perfect DRACH matches, comparable to the 96% and 95% observed for infrequent and background sites, respectively (Fig. 3e; Supplementary Table 4). These results indicate that TC sites have the same canonical sequence preference associated with the MTC as other infrequent or background sites have.

**Fig. 3.**
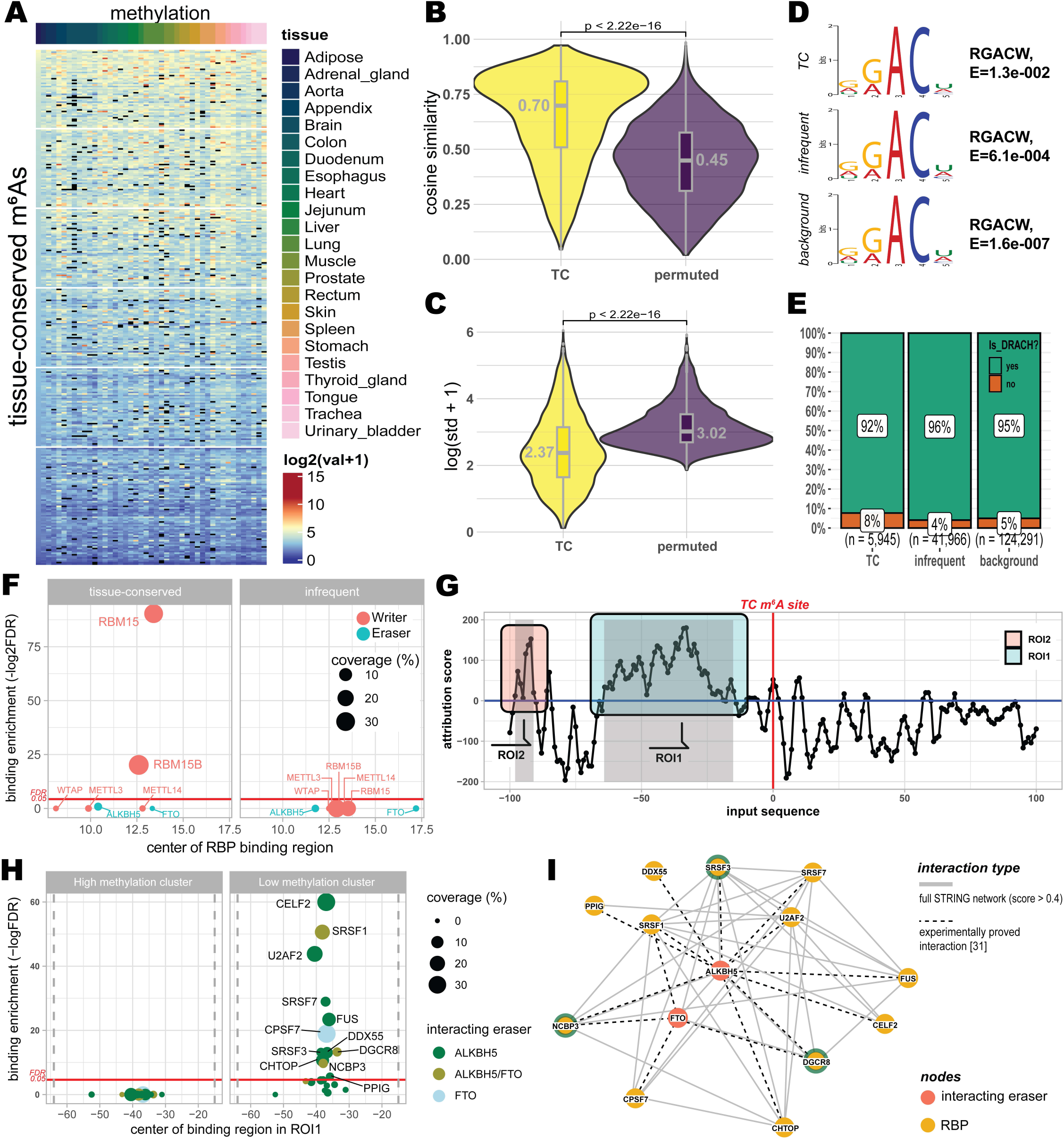
Regulatory mechanisms and cis-encoded features of TC m6A sites. (A) Methylation levels of TC m6A sites across 24 human tissue types. (B and C) Comparison of cosine similarity and standard deviation of methylation levels between TC sites and a random model. Statistical significance was determined by a two-sided Wilcoxon rank-sum test (P<2.22*10-16). (D) De novo motif analysis using MEME Suite identifying the 5-RGACW-3’ consensus sequence across TC, infrequent, and background site classes. (E) Percentage of m6A sites containing perfect matches to the canonical DRACH motif across the three site classes. (F) Enrichment of m6A writer and eraser binding near TC and infrequent sites relative to background sites. Significant enrichment is shown for RBM15 and RBM15B near TC sites. (G) Attribution score of regressing sequence using the fine-tuned m6A-BERT model. Two key regions of interest (ROIs) strong contribution (i.e., positive attribution score) to regressing the methylation levels were identified. Here, a region where positive attribution score continued more than 5 bp was defined as ROI for analysis: ROH (−64 to −15 bp) and ROI2 (−98 to −91 bp). (H) Enrichment of RNA-binding proteins (RBPs) within ROH. Colors indicate RBPs that have

To investigate whether the deposition of TC site involves specific accessory proteins, we analyzed binding profiles of 217 RNA-binding proteins (RBPs), including 5 writers, 2 erasers, and 18 readers (WERs), using the POSTAR3 database^31^. We first assessed the enrichment of writer and eraser binding near TC and infrequent sites relative to background sites. Among all writers, only *RBM15* (FDR = 8.4 × 10⁻⁷) and *RBM15B* (FDR = 6.4 × 10⁻²⁸) showed significant enrichment near TC sites (Fig. 3f). Their binding centers were located 13.4 bp (*RBM15*) and 12.6 bp (*RBM15B*) from TC sites on average, indicating direct proximity (Supplementary Fig. 12). In contrast, these proteins did not show enrichment near infrequent sites (Fig. 3f). Expression analysis revealed that both *RBM15* and *RBM15B* were stably expressed across all 24 tissues (Supplementary Fig. 13), further supporting their role in tissue-independent methylation. While 25 additional RBPs that are not known m^6^A writer/eraser were also consistently expressed and enriched near TC sites (Supplementary Figs. 13 and 14), protein-protein interaction analysis using STRING^32^ showed that only *RBM15* and *RBM15B* directly interact with *METTL3* or *METTL14* (confidence score > 0.9; Supplementary Fig. 15). These findings imply *RBM15* and *RBM15B* as candidate accessory proteins associated with TC sites and compatible with a role in facilitating MTC engagement at these sites.

Although TC sites are reproducibly methylated across tissues, their methylation levels in a tissue substantially vary among them (Fig. 3a), suggesting that additional factors may modulate methylation stoichiometry. A previous study in yeast has shown that cis-sequence features can influence m^6^A stoichiometry ^5^. Motivated by this, we fine-tuned m^6^A-BERT^27^ to regress average methylation levels of TC sites using ±100 bp flanking sequences. The model achieved an *R²* = 0.70, indicating that local sequence context accounts for a substantial portion of the variation in methylation levels across TC sites. To identify specific sequence regions contributing to this prediction, we applied the integrated gradient analysis to the fine-tuned model. This revealed two upstream regions of interest (ROIs; see Methods; Fig. 3g) with strong attribution scores: ROI1 (−64 to −15 bp) and ROI2 (−98 to −91 bp). These regions may contain cis-elements associated with co-factors modulating m^6^A deposition or removal. To explore potential co-factors, we stratified TC sites into high- and low-methylation clusters based on their average methylation levels (Supplementary Fig. 16) and examined RBP binding enrichment between high-methylation cluster and low-methylation cluster within ROI1 sequences. To focus on tissue-independent effect, we restricted analysis to184 RBPs in POSTAR3 database^31^ that maintained relatively consistent expression across tissues. No RBP was significantly enriched in the high-methylation cluster (Fig. 3h). In contrast, 85 RBPs showed significant enrichment in the low-methylation cluster (Fisher’s exact test, FDR < 0.01; Supplementary Fig. 17), suggesting a potential association with decreasing methylation levels. Among these RBPs, only 12 have been reported to physically interact with the m^6^A erasers ALKBH5 or FTO (Fig. 3h), including *DGCR8*, *NCBP3*, and *SRSF3*^33^. Integrating STRING^32^ interaction with published experimental protein interactome analysis results^33^ further confirmed these interactions (Fig. 3i), where DGCR8 uniquely interacting with both erasers. These observations raise the possibility that a subset of upstream-binding RBPs may be associated with demethylase recruitment towards TC sites in low-methylation clusters. A similar enrichment pattern was observed for ROI2 in the low-methylation but not in the high-methylation cluster (Supplementary Fig. 18), further supporting a potential link between upstream RBP binding and modulation of methylation stoichiometry. In summary, we identify *RBM15* and *RBM15B* as the only writer-associated proteins significantly enriched near TC sites, potentially functioning as accessory factors linking the MTC to these sites. In parallel, differences in TC sites methylation levels are linked to upstream binding of additional RBPs, particularly within ROI1 and ROI2, including proteins reported to interact with the erasers *ALKBH5* or *FTO*. Together, these results support a layered model in which factors near the site and upstream sequence-associated RBPs may contribute to TC m^6^A site deposition and site-specific modulation of methylation stoichiometry.

### Tissue-conserved m^6^A sites are associated with YTHDF/UPF1-mediated mRNA decay

m^6^A controls gene expression through the recruitment of reader proteins. To examine whether TC m^6^A sites are associated with conserved expression regulation, we first examined expression patterns of TC site-containing genes. Across tissues, these genes showed broadly similar expression profiles (Fig. 4a), with significantly higher cosine similarity and lower standard deviation than the expected under a null distribution (Wilcoxon rank-sum test, *P* < 2.22 × 10⁻¹⁶; Fig. 4b,c). We further observed a significant negative correlation between TC-site methylation levels and the expression levels of their host genes (Spearman’s ρ = −0.50, *P* < 2.2 × 10⁻¹⁶; Supplementary Fig. 19), indicating that higher TC methylation is associated with lower gene expression.

**Fig. 4.**
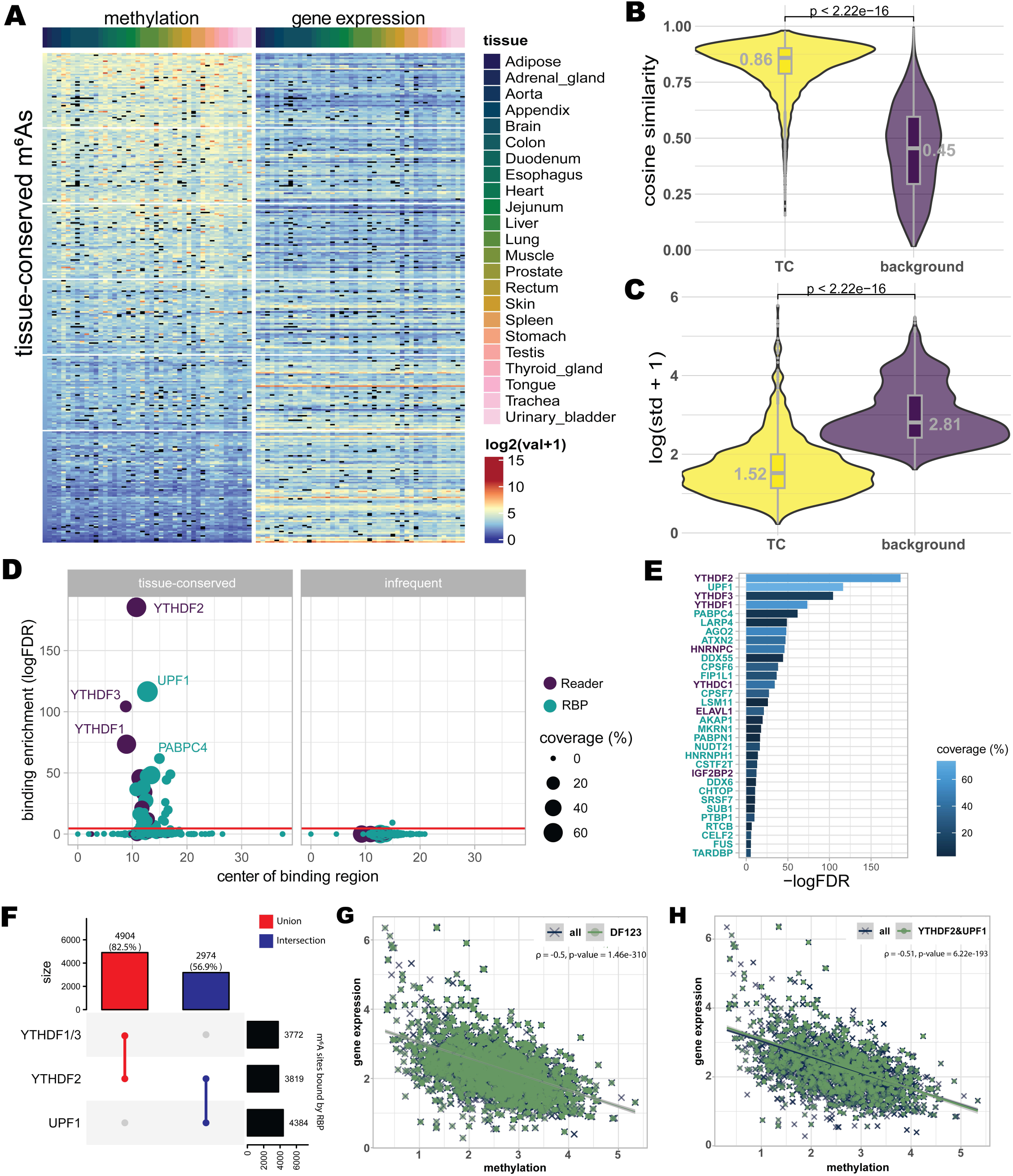
TC m6A sites mediate context-independent gene repression through m6A-mediated decay. (A) Methylation levels of TC sites and expression levels of associated genes containing TC m6A sites across 24 human tissues. (B and C) Comparison of cosine similarity and standard deviation of gene expression levels harboring TC sites and background genes. Statistical significance was determined by a two-sided Wilcoxon rank-sum test (P<2.22×10-16). (D) Enrichment of RBP binding near TC sites, analyzed using POSTAR3 data. The top four enriched proteins (YTHDF2, UPF1, YTHDF3, and YTHDF1) are highlighted, with mean distances from their binding centers to TC sites indicated. Red line represents the significance threshold (FDR = 0.05). (E) Enrichment scores (FDR) for the top RBPs associated with TC sites, including the YTHDF family and UPF1. (F) UpSet plot showing the overlap of TC sites bound by YTHDF1, YTHDF2, YTHDF3, and UPF1. In total, 82.5% of TC sites are bound by at least one YTHDF protein, and 56.9% are co-bound by YTHDF2 and UPF1. (G) Correlation between TC site methylation and host gene expression for sites bound by YTHDF1-3. (H) Correlation between TC site methylation and host gene expression for sites co-bound by both YTHDF2 and UPF1.

To identify the readers and decay-associated factors likely responsible for TC site-linked transcript repression, we analyzed binding enrichment of RBPs near TC and infrequent sites relative to the background sites, excluding writers and erasers. Only 32 RBPs were significantly enriched near TC sites (FDR < 0.05; Fig. 4d), including 7 canonical m^6^A readers and 25 additional RBPs, indicating that TC sites are associated with a selective subset of RBPs. Among these, *YTHDF2* shows the strongest enrichment (FDR = 2.99 × 10⁻⁸¹), implicating this principal m^6^A-decay reader as a major mediator of TC-site transcript downregulation. In addition, *YTHDF3* and *YTHDF1*, known to act redundantly with YTHDF2 to promote mRNA degradation^34^, ranked third and fourth (FDR = 4.91 × 10⁻⁴⁶ and 1.23 × 10⁻³², respectively; Fig. 4d,e). Consistent with these rankings, 82.5% of TC sites overlapped binding regions of at least one of *YTHDF1-3* (Fig. 4f), and TC sites bound by *YTHDF1-3* displayed a negative association between methylation level and host-gene expression (Spearman’s ρ = −0.5, *P* = 1.46 × 10⁻^310^; Fig. 4g). Together, these results indicate that TC sites are preferentially engaged by the YTHDF family and that their methylation is associated with lower transcript abundance.

Among the remaining enriched RBPs, *UPF1*, a factor classically associated with nonsense-mediated decay, was the second most enriched factor near TC sites (FDR = 2.73 × 10⁻⁵¹, Fig. 4d,e). Although *UPF1* is not typically considered as a core m^6^A reader, recent work has shown that *UPF1* can cooperate with *YTHDF2* to promote degradation of m^6^A-methylated mRNAs ^35^. In our data, over half of TC sites (56.9%) were co-bound by both *YTHDF2* and *UPF1* (Fig. 4f), and these co-bound sites exhibited even stronger inverse methylation-expression relationship (ρ = −0.51, *P* = 6.22 × 10⁻¹⁹³; Fig. 4h) than YTHDF1-3 co-bound (Fig. 4g). Moreover, binding peak centers of *YTHDF2*, *UPF1*, *YTHDF3*, and *YTHDF1* were located in close proximity to TC sites (10.7 bp, 12.8 bp, 8.8 bp, and 8.9 bp away, respectively; Fig. 4d), consistent with direct recognition of TC-methylated transcripts. Notably, these decay-associated RBPs were not enriched near infrequent sites, underscoring the specificity of this reader-decay engagement with TC sites. Collectively, these findings support a conserved TC m^6^A decay program centered on YTHDF1-3 and UPF1 that links stable TC methylation to transcript repression across tissues.

### TC m^6^A genes maintain expression homeostasis in essential cellular pathways

To characterize the biological function of these 1,386 stably expressed and methylated TC m^6^A genes, we performed Gene Ontology (GO) enrichment analysis. These genes converged on three broad functional categories (Fig. 5a). The first category relates to cellular quality control and homeostasis, including macroautophagy, ERAD pathway, regulation of protein catabolic processes, and ribosome biogenesis, whose functions involved in degrading damaged components and maintaining protein integrity. The second involves intracellular organization and inter-organelle coordination functions, such as Golgi vesicle transport, nuclear transport, RNA splicing, and inner mitochondrial membrane organization, reflecting roles in molecular trafficking between compartments. The third includes stress responses and host-pathogen interaction processes, including viral life cycle, viral process, stem cell population maintenance, and cellular response to unfolded protein stress, suggesting roles in adaptation to environmental and pathogenic challenges. Together, these results indicate that TC m^6^A genes are enriched in core cellular processes that underpin homeostasis, intracellular logistics, and stress responsiveness.

**Fig. 5.**
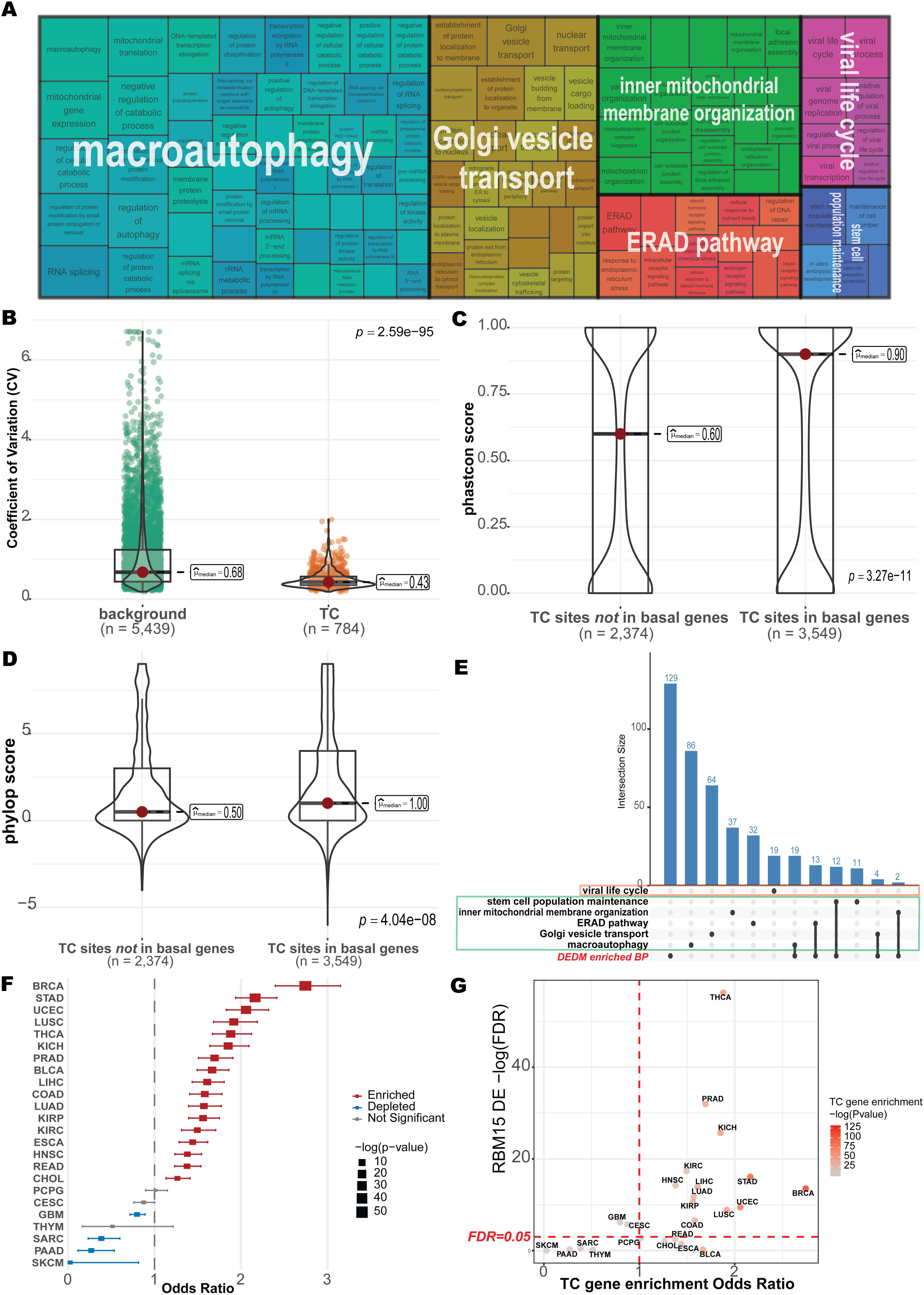
TC m6A methylation stabilizes essential housekeeping and preparedness pathways to maintain cellular homeostasis. (A) Gene Ontology (GO) enrichment analysis for 1,386 genes containing TC m6A sites. Key enriched biological process (BP) terms are grouped into functional categories (B) Comparison of the coefficient of expression variation across 24 tissues forTC m6A genes involved in basal preparedness and control gene sets. (C and D) Evolutionary conservation of TC m6A sites in basal preparedness TC genes compared to other TC m6A sites. (E) UpSet plot of enriched GO:BP terms. DEDM enriched BP represents enriched terms using 60 TC m6A genes that were also identified as differentially expressed and differentially methylated (DEDM) during Herpes Simplex Virus 1 (HSV-1) infection. Enriched terms are primarily related to cellular logistics and homeostasis, not canonical immune responses. (F) Pan-cancer enrichment analysis of the tissue-conserved (TC) m6A gene set (n = 1,386 genes) across 24 cancer types in TCGA. Odds ratios (OR) represent the likelihood of TC m6A genes being differentially expressed compared to other differentially expressed genes. (G) Differential expression enrichment odd ratio of TC m6Agene set (n = 1,386 genes) and the differential expression significance (FDR) of RBM15B across 24 TCGA cancers.

Given that these TC m^6^A genes are consistently expressed across tissues and enriched in pathways requiring sustained basal activity, we hypothesized that TC m^6^A methylation may help establish gene-specific basal expression set-points. To test this, we examined the expression variability of TC m^6^A genes involved in cellular quality control and basal stress preparedness. These genes exhibited significantly lower coefficients of variation across 24 tissues compared to background genes (median coefficient of variance = 0.43 vs. 0.68; Wilcoxon rank-sum test, *P* = 2.59 × 10⁻⁹⁵; Fig. 5b). This reduced variability suggests that TC m^6^A methylation contributes to tighter expression buffering across tissues. To assess the evolutionary significance of this constraint, we compared the sequence conservation of TC sites located within these basal preparedness genes to TC sites outside them. TC sites in basal genes displayed significantly higher phastCons scores (median 0.90 vs. 0.60; Wilcoxon rank-sum test, *P* = 3.27 × 10⁻¹¹; Fig. 5c) and phyloP scores (median 1.00 vs. 0.50; Wilcoxon rank-sum test, *P* = 4.04 × 10⁻⁸; Fig. 5d), indicating stronger selective pressure to preserve TC m^6^A sites within these functionally essential genes. Collectively, these findings suggest that TC m^6^A methylation provides a conserved mechanism for stabilizing basal expression levels in genes essential for cellular quality control and stress preparedness.

To evaluate the potential role of TC m^6^A genes in pathogen response, we intersected our gene set with genes reported as differentially expressed and differentially m^6^A-methylated (DEDM) during Herpes Simplex Virus 1 (HSV-1) infection^4^. We identified 60 overlapping genes (e.g., *VCP, FOXO3, KEAP1, MYC, EXOC7, SUMO1*; Supplementary Table 5), many of which function in vesicle trafficking (*EXOC7, BET1L*)^36–38^, ER-associated degradation (*VCP, TRAPPC10*)^39,40^, and mitochondrial remodeling (*SCO2, ETF1*)^41,42^. Functional enrichment analysis using these genes yielded 179 GO biological process terms. An UpSet analysis revealed that these enriched overlapping genes intersected predominantly with macroautophagy, Golgi vesicle transport, the ERAD pathway, and mitochondrial organization categories, but notably not with immune-response categories (Fig. 5a, e), consistent with previous reports that HSV-1 reprograms m^6^A primarily to rewire host logistics rather than broadly alter immune gene regulation^4^. Several of the overlapping TC genes illustrate this pattern: *VCP* supports proteostasis remodeling during viral protein overload^40,43^, *SUMO1* and *DAXX* participate in chromatin silencing and reactivation of viral genes^44,45^, and *USP22/UBE2L3* contribute to ubiquitin cycling required for viral assembly and egress^46,47^. These findings suggest that HSV-1 exploits TC m^6^A-regulated housekeeping pathway to support replication, positioning TC m^6^A methylation not only as a contributor to cellular homeostasis but also as a potential point of vulnerability to viral exploitation.

Having established that TC m^6^A methylation contributes to expression homeostasis in essential cellular pathways, we next asked whether this regulatory layer is preferentially disrupted in cancer. We performed differential expression (DE) analysis across 24 cancer types in The Cancer Genome Atlas (TCGA), comparing primary tumors to matched normal samples. For each cancer type, we used Fisher’s exact test to assess whether TC m^6^A genes were more likely to be differentially expressed than expected given the background transcriptome. TC m^6^A genes were significantly enriched among DE genes in 17 of 24 cancer types (Fisher’s exact test, *OR* > 1, FDR < 0.05; Fig. 5f), with the strongest effects observed in breast invasive carcinoma (BRCA; OR = 2.74) and stomach adenocarcinoma (STAD; OR = 2.16). In seven types of cancer (SKCM, PAAD, SARC, THYM, GBM, CESC, and PCPG), TC m^6^A genes showed no DE enrichment (Fisher’s exact test, OR < 1 or FDR > 0.05), suggesting that their expression remains relatively stable in these contexts. Overall, these results indicate that TC m^6^A genes are disproportionately subject to transcriptional alteration across most cancer types examined.

Given our earlier identification of *RBM15* and *RBM15B* as candidate writers linked to TC m^6^A deposition, we examined whether these regulators were themselves dysregulated in the same cancers. *RBM15* was significantly differentially expressed in 15 of 24 types of cancer (62.5%), and *RBM15B* in 7 of 24 (Supplementary Fig. 20). In all seven cancers where *RBM15B* was dysregulated, *RBM15* was also significantly altered, suggesting coordinated disruption of both regulators. Across cancer types, the degree of TC gene enrichment among DE genes was broadly correlated with the significance of *RBM15/B* dysregulation. Cancer types with strong TC gene enrichment (high odds ratio) generally also showed significant *RBM15/B* dysregulation (FDR < 0.05), as seen for BRCA, STAD, THCA, PRAD, and KICH in the upper-right quadrant (Fig. 5g). Conversely, cancers where TC m^6^A genes were not enriched among DE genes (e.g., SKCM, PAAD, SARC, and THYM) also showed no significant *RBM15/B* alteration, clustering in the lower-left quadrant (Fig. 5g). However, two cancers, including BLCA and ESCA, showed elevated TC gene enrichment without significant *RBM15/B* dysregulation, suggesting that TC gene disruption in these cancers may occur through alternative mechanisms. This overall pattern of concordance between writer dysregulation and TC m^6^A gene disruption across most cancer types is consistent with a model in which altered *RBM15/B* expression contributes to the transcriptional changes observed at TC m^6^A genes.

## Discussions

By integrating 67 MeRIP-seq datasets across 24 normal human tissues, we identified 5,945 tissue-conserved (TC) m^6^A sites with consistent methylation regardless of tissue origin, donor background, or experimental protocol. Sites were validated using single-base-resolution annotation, permutation testing, and cross-dataset filtering, and further supported by detection across diverse cell lines and conditions. Recovery of constitutively methylated genes (*ACTB*, *BSG*, and *MALAT1*) adds additional confidence, pointing to a core m^6^A program that operates independently of tissue identity.

TC sites differ from infrequent and background sites by molecular and evolutionary features, including stronger enrichment near stop codons, elevated methylation levels across transcript regions, greater spatial clusters, and higher sequence conservation. Their positioning also aligns with current models in which exon junction complex mediates suppression of m^6^A deposition^24–26^. Infrequent sites tend to occupy long internal exons, suggesting dependency on locally permissive architecture. A fine-tuned m^6^A-BERT accurately classifies TC sites, further supporting distinct local cis-regulatory features that support stable methylation.

*RBM15* and *RBM15B* emerge as candidate accessory factors for TC m^6^A deposition: they are the only writer-associated proteins enriched near TC sites, and both are stably expressed across tissues and known to interact with *METTL3/METTL14*, consistent with a model in which *RBM15/B* help recruit the methyltransferase complex to TC sites. Despite consistent deposition, TC sites vary in methylation levels among themselves in a tissue. A regression model (*R*^2^ = 0.70) revealed that local flanking sequences predict methylation levels, and integrated gradient analysis identified two upstream regions. RBPs enriched at low-methylation TC sites within these regions include factors known to interact with the erasers *ALKBH5* and *FTO*, suggesting that upstream RBP binding may recruit demethylases and reduce methylation. This adds an important level of nuance: even stable sites may be subject to sequence-encoded dampening to prevent over-methylation. These findings suggest that the cell maintains TC sites not just through recruitment, but through an active and balanced competition between writers and erasers determined by the flanking sequence. Together, the data support a layered architecture, in which *RBM15/B* help establish baseline TC methylation, while upstream cis-elements and associated RBPs fine-tune stoichiometry.

Functionally, TC sites are associated with transcript downregulation. Genes harboring TC sites show reduced expression variability across tissues, and TC-site methylation inversely correlates with transcript abundance. TC sites are preferentially bound by a selective subset of RBPs; YTHDF2 shows the strongest enrichment, with YTHDF1/3 acting redundantly in degradation. UPF1, co-bound with YTHDF2, strengthens the inverse methylation-expression relationship. The lack of enrichment at infrequent sites underscores specificity. These findings support a conserved mRNA-decay program centered on YTHDF1-3 and UPF1.

The functional TC m^6^A-mediated decay pathway suggests that this stable layer serves as a “buffering” mechanism for cellular homeostasis. The 1,386 genes marked by TC sites are enriched in core cellular processes such as autophagy, protein quality control and intracellular trafficking, forming a backbone for cellular homeostasis rather than tissue-specific identity programs. Their reduced expression variability and stronger evolutionary constraint at TC sites is consistent with TC m^6^A stabilizing basal expression of essential housekeeping pathways. TC m^6^A sites may prevent toxic transcript accumulation and ensure rapid turnover, allowing these essential systems to remain primed and responsive by maintaining a constant degradation for these genes. This architecture may also create entry points during stress and infection: in HSV-1 datasets^4^, TC-marked genes intersect primarily with logistics and housekeeping, not broad immune pathways, suggesting viral exploitation of pre-existing homeostatic circuitry. In the cancer context, TC m^6^A genes are disproportionately differentially expressed across many TCGA tumor types, often coinciding with dysregulation of *RBM15/B*, consistent with perturbation of this stable layer contributing to transcriptional remodeling of essential programs.

Several limitations of this study should be noted. Our identification relies on MeRIP-seq, which provides regional rather than single-nucleotide resolution, and integration with single-base resolution references only partially mitigates this. Sample coverage for certain tissues is limited, and the focus on shared sites may underrepresent tissue-specific m^6^A. Additionally, while our RBP enrichment analysis is compelling, direct perturbation experiments will be essential to confirm the predicted functional roles of *RBM15/B* and upstream RBPs in TC m^6^A regulation. In cancer, the lack of matched tumor-normal m^6^A profiling prevents direct attribution of differential expressions to altered TC-site methylation, a question that expanded epitranscriptomic datasets will help resolve.

In conclusion, we identify a consistently methylated subset of the human m^6^A landscape with distinctive sequence context, regulatory associations, and functional coupling to mRNA decay. This tissue-conserved layer appears to stabilize essential gene regulation and constitutes a coherent epitranscriptomic program that can be disrupted in disease.

## Methods

### Data resources and pre-processing

Human tissue MeRIP-seq datasets were retrieved from the Chinese Brain Bank Center^9^ (GSE122744) and the National Genomics Data Center^10^ (CRA001315), spanning 24 normal tissue types. Quality control was performed using fastp^48^ and adapter sequences were trimmed using cutadapt^49^ (v2.10; --minimum-length 25 --pair-filter=any). Reads were aligned to the GRCh38 genome assembly using HISAT2^50^ (v2.2.1; --no-mixed --no-discordant). Transcriptome quantification was conducted using the GENCODE^51^ gene annotation (release 38) and featureCounts^52^ (v2.0.1; -F GTF -t exon -g gene_id -s 2 -p -B -M --fraction --ignoreDup -T).

### Identification of high-confidence m^6^A sites

To establish a robust reference set of m^6^A sites, we integrated 427,760 sites from m^6^A-Atlas v2.0^11^ and 148,655 sites identified via the GLORI method^12^, totaling 480,636 unique m^6^A sites across 30,227 genes. To mitigate potential biases associated with antibody dependency or low sequencing coverage, we applied the following stringency filters: (i) Genomic location of m^6^A sites was restricted to exonic regions; intronic, intergenic, and distal downstream sites were excluded to focus on the processed mRNA landscape. (ii) Sites were required to be detected by at least two distinct sequencing technologies. To ensure independence, miCLIP^53^ and miCLIP2^53^ were treated as a single methodological category. This refined set was utilized as the background m^6^A landscape for all subsequent analyses.

### Identification of tissue-conserved (TC) m^6^A sites

m^6^A peaks were predicted by exomePeak2^13,14^. High-quality peaks were selected based on the following criteria: (i) FDR < 0.05; (ii) assignment to a single gene with detectable expression; (iii) non-zero peak read counts in both IP and INPUT samples; and (iv) a peak length of < 1,000 bp. The high-confidence m^6^A sites identified in the previous step were intersected with these m^6^A peaks to assess their distribution across tissues. To account for the technical variations in sequencing depth and sample quality, the CBBC and NGDC datasets were analyzed independently. For each dataset, a permutation-based null distribution was generated by shuffling m^6^A site occurrences across samples to model random methylation. Observed site frequencies were compared against this background distribution, and P-values were calculated for each site and adjusted using the Benjamini–Yekutieli (BY) method^54^. Significantly shared sites (*FDR* < 0.01) were required to be detected across all tissue types within their respective datasets (7 tissues in CBBC and 23 tissues in NGDC). Tissue-conserved (TC) m^6^A sites were then defined as those consistently identified as significantly shared across all 24 tissue types represented in both datasets. For comparison, infrequent sites were defined as those detected in three or fewer tissue types.

### Molecular and evolutionary feature analysis

The spatial distribution of m^6^A sites along transcript architecture was analyzed using the MetaTX R package^55^. To ensure statistical comparability and avoid bias due to varying set sizes, the infrequent and background m^6^A site sets were randomly downsampled to match the total number of identified TC sites (*n* = 5,945) before metagene plotting. Evolutionary conservation scores were retrieved using the GenomicScores R package^56^. Specifically, we utilized the phyloP100way.UCSC.hg38 and phastCons100way.UCSC.hg38 data packages, which provide conservation scores based on the 100-way vertebrate alignment to the GRCh38 genome. Scores were extracted for the focal adenosine of each m^6^A site and compared across site categories. Intra-class clustering was assessed by calculating the frequency of neighboring m^6^A sites within specific genomic windows of ±25, 50, 100, and 200 bp. For a given site, neighbors were only counted if they belonged to the same category (TC, infrequent, or background) on the same transcript. Additionally, we assessed the enrichment of previously reported “clustered m^6^A sites” from the GLORI dataset^12^ within each category using Fisher’s exact test.

### Sequence-based deep learning with m^6^A-BERT

#### TC m^6^A site classification

To distinguish between tissue-conserved and infrequent methylation, we utilized 5,945 TC m^6^A sites as the positive set and a randomly downsampled subset of 5,945 infrequent sites (from a pool of 41,966) as the negative set. For each site, the genomic sequence was extended by ±50 bp, resulting in 101 bp input sequences represented by DNA-encoded nucleotides (A, T, G, C). Classification was performed by fine-tuning the m^6^A-BERT model^27^, which was pre-trained on 427,760 human m^6^A sites from m^6^A-Atlas v2.0^11^. Following the BERT architecture, input sequences were represented as *k*-mer tokens generated with a sliding window of length *k* = 3. To maintain input consistency, special tokens ([CLS] and [SEP]) were added, and sequences were padded ([PAD]) to a maximum length of 120 tokens. A binary classification layer was appended to the pre-trained m^6^A-BERT base to predict TC (1) versus infrequent (0) status. The dataset was partitioned into training (70%), testing (20%), and validation (10%) sets. The model was fine-tuned over 50 epochs with a learning rate of 1 × 10^−6^ and the final model (*k* = 3) was selected based on the maximum Area Under the ROC Curve (AUC) achieved on the validation set. **TC m^6^A methylation regression:** To quantify the contribution of local sequence context to TC site methylation levels, we trained a regression model using the 5,945 TC m^6^A sites. For each site, the target variable was defined as the mean methylation level across all NGDC tissue samples. To prevent the model from over-relying on the invariant m^6^A consensus motif, the central adenosine and its ±2 bp flanking regions were replaced with random nucleotides after sequence extraction (±50 bp). We employed the same pre-trained m^6^A-BERT model with an additional linear regression layer. The dataset was partitioned into training (70%), testing (20%), and validation (10%) sets. Fine-tuning was conducted over 50 epochs with a learning rate of 1 × 10^−6^. The final model was selected based on the highest coefficient of determination (*R²*) achieved on the validation set. **Model interpretation:** To identify the sequence elements driving the model’s predictions, we employed a gradient-based attribution method^57^. The attribution score *IG* for an input token *x* relative to a reference token *x_ref_* was calculated as:

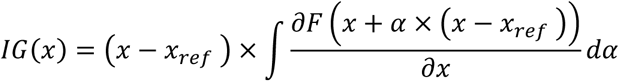

The reference input *x_ref_* consisted of a sequence of [PAD] tokens, anchored by [CLS] and [SEP] tokens. Attributions were calculated at the embedding layer using the Captum library^58^. Nucleotide-level importance scores were derived by aggregating and averaging attribution scores across all corresponding tokens, weighted by their frequency. Regions of Interest (ROI) were defined as contiguous genomic segments of positive attribution scores > 5 bp in length, with adjacent segments merged if the non-positive score gap was ≤ 3 bp.

### RBP binding and enrichment analysis

To identify RBPs potentially regulating TC m^6^A sites, we retrieved cross-linking immunoprecipitation followed by sequencing (CLIP-seq) data from the POSTAR3 database^31^. RBP binding peaks > 100 bp were excluded to ensure high-resolution mapping. We implemented a filtering pipeline using zFPKM values^59,60^ to identify active genes (median *zFPKM* > −3) across the 24 tissues. RBPs were prioritized if they exhibited consistent expression and lacked significant differential expression across tissues. Enrichment was determined using a one-sided Fisher’s exact test (alternative = “greater”), comparing binding frequencies between TC/infrequent sites and between high/low methylation clusters. Nominal *P*-values were adjusted using the Benjamini–Hochberg (BH) method^61^. RBPs enriched in the low methylation cluster at ROI1 were further analyzed to assess interaction with erasers using the experimental proteomics data^33^ (Limma adjusted *P*-value < 0.05; SAINT score > 0.74) and STRING database (v11.5; full STRING network with confidence score > 0.4)^32^.

### Functional and transcriptomic analysis

For each TC and infrequent m^6^A site, the methylation level was defined by the enrichment ratio of the overlapping peak. This ratio was calculated as the normalized IP read count divided by the normalized input read count. Gene expression levels (FPKM) were quantified using featureCounts as detailed in the “Data resources and pre-processing” section. The correlation between methylation levels and corresponding transcript abundance was evaluated using Spearman’s rank correlation. To assess expression stability, the coefficient of variation (CV) for gene expression was calculated across the 24 normal tissues and compared between TC m^6^A genes and background gene sets using the Wilcoxon rank-sum test. To characterize the biological roles of genes containing TC m^6^A sites, GO enrichment analysis was performed using the clusterProfiler R package^62,63^ (v4.14.6). To reduce redundancy and visualize the functional landscape, semantic similarity analysis of the enriched GO terms was conducted using the rrvgo R package^64^ (v1.18.0). For the pan-cancer analysis, we retrieved transcriptomic data for 24 cancer types from TCGA using the TCGAbiolinks R package^65^. Our analysis focused on paired primary tumor and solid tissue normal samples to evaluate the differential regulation of TC m^6^A-harboring genes in a clinical context. For each TCGA cohort, differential gene expression between tumor and normal tissues was performed using the DESeq2 R package^66^ (FDR < 0.05, |log₂(Fold Change)| > 1). To determine if TC m^6^A genes were disproportionately altered, a Fisher’s exact test was applied for each cancer type, comparing the frequency of differential expression among TC m^6^A genes against the background transcriptome. Finally, to assess the concordance with writer dysregulation, the enrichment odds ratios of TC m^6^A genes were compared against the differential expression significance (FDR) of *RBM15* and *RBM15B* across the 24 cancer types.

### Statistical Analysis

All statistical analyses were performed using R (v4.4.1). Unless otherwise specified, all statistical tests were two-sided, and a *P*-value (or *FDR*) < 0.05 was considered statistically significant. For comparisons between two groups of continuous variables (e.g., conservation scores, *CV*, or methylation levels), the Wilcoxon rank-sum test was employed. Enrichment of categorical variables (e.g., RBP binding or site overlap) was evaluated using Fisher’s exact test. Correlations were calculated using either Pearson’s *r* (for linear relationships in model training) or Spearman’s rank correlation (*ρ*) (for non-linear transcriptomic associations). Multiple testing corrections were performed using the BH or BY procedure to control the false discovery rate. For machine learning tasks, datasets were partitioned into non-overlapping training (70%), testing (20%), and validation (10%) sets to prevent data leakage and ensure model generalizability. Random downsampling was used for all comparative analyses between TC and infrequent sites to eliminate bias arising from unequal sample sizes. No data points were excluded from the analysis unless they failed the initial quality control criteria (e.g., low read counts or excessive peak length) described in the Methods.

## Data and code availability

The code used for all primary analyses and corresponding data are available on GitHub at https://github.com/Huang-AI4Medicine-Lab/TC-m6A.

## Acknowledgement

We thank members of Drs. Shou-Jiang Gao and Yufei Huang laboratories for technical assistance and discussions.

## Author contribution

Conceptualization, Y.H.; methodology, S.J., T.Z.; investigation, S.J., T.Z., S.-J.G., Y.H.; writing – original draft, S.J, Y.H.; writing – review & editing, S.J., T.Z., S.-J.G., Y.H.; funding acquisition, S.-J.G., Y.H.; supervision, S.-J.G., Y.H.

