## SupplementaryFigures for "Systematic identification of tissue-conserved m^6^A sites reveals a stable epitranscriptomic regulatory layer controlling essential genes"

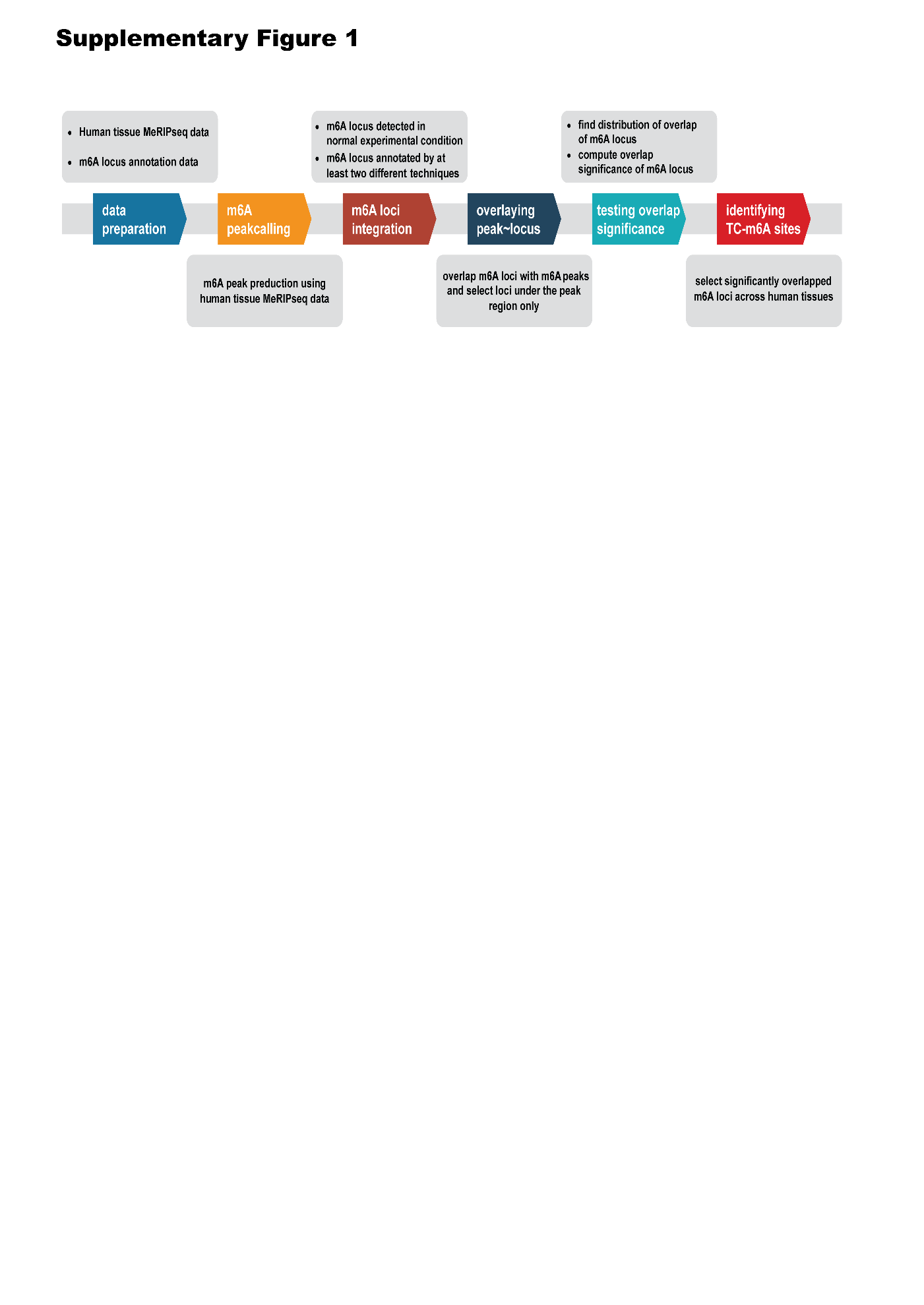

Supplementary Fig. 1 | A bioinformatics pipeline that integrates MeRIP-seq data with high-confidence single-nucleotide m6A annotations.

Frist human tissue MeRIP-seq data and single-nucleotide base m6a annotation data were collected. Second, we called the m6A peaks using exomePeak2 R package against human tissue MeRIP-seq data. Third, we rigorously selected high confidence m6A sites annotations, which met two criteria. Fourth, we overlayed the predicted peaks and high confidence m6A sites to further refine the sites under the peak region only. Fifth, we compute the overlap distribution of those m6A sites and test the overlap significance of individual m6A site. Finally, we identified significantly overlapped m6A sites across human tissues and determined tissue-conserved m6A sites.

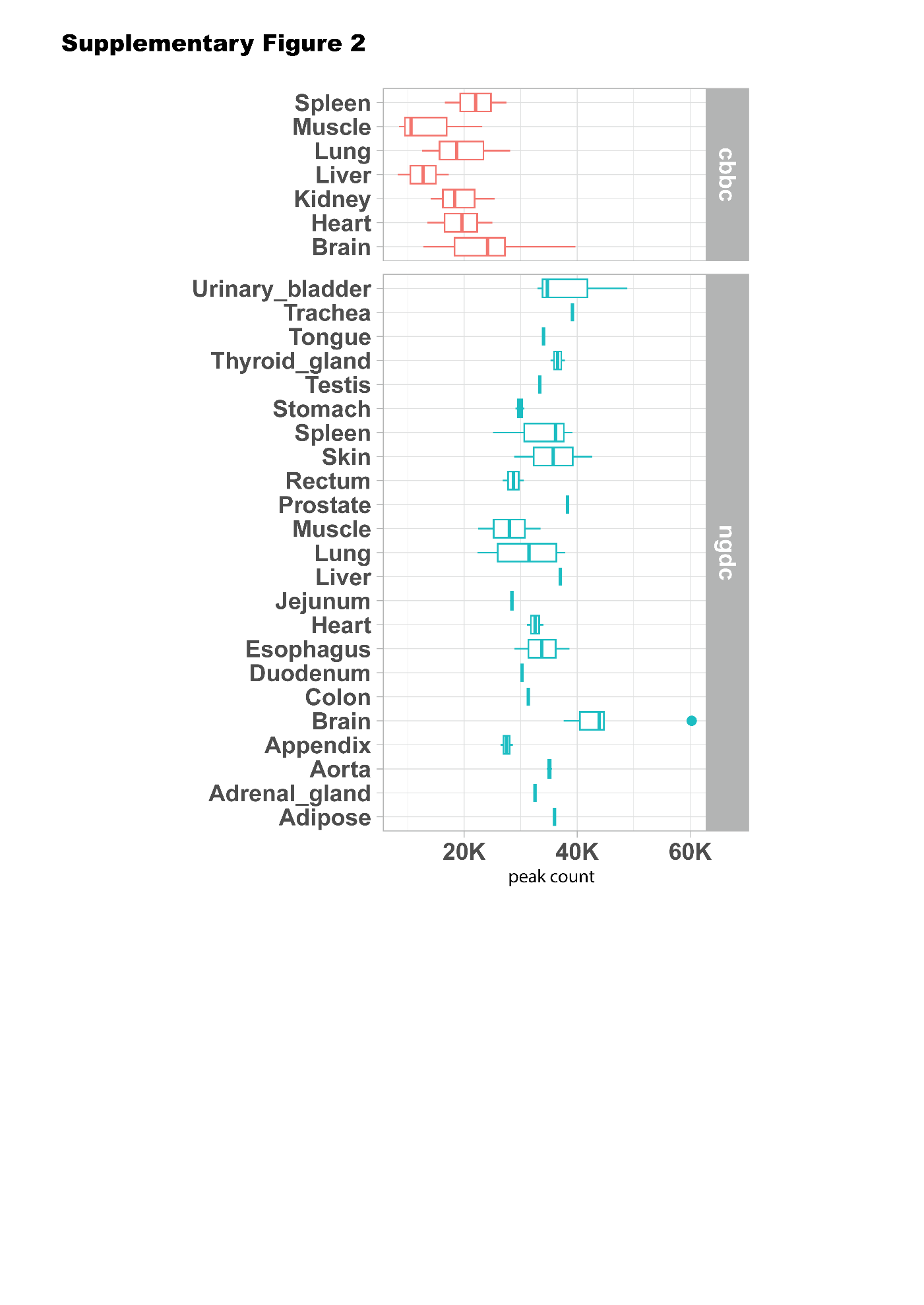

Supplementary Fig. 2 | Average number of m6A peaks in each tissue type in each dataset respectively.

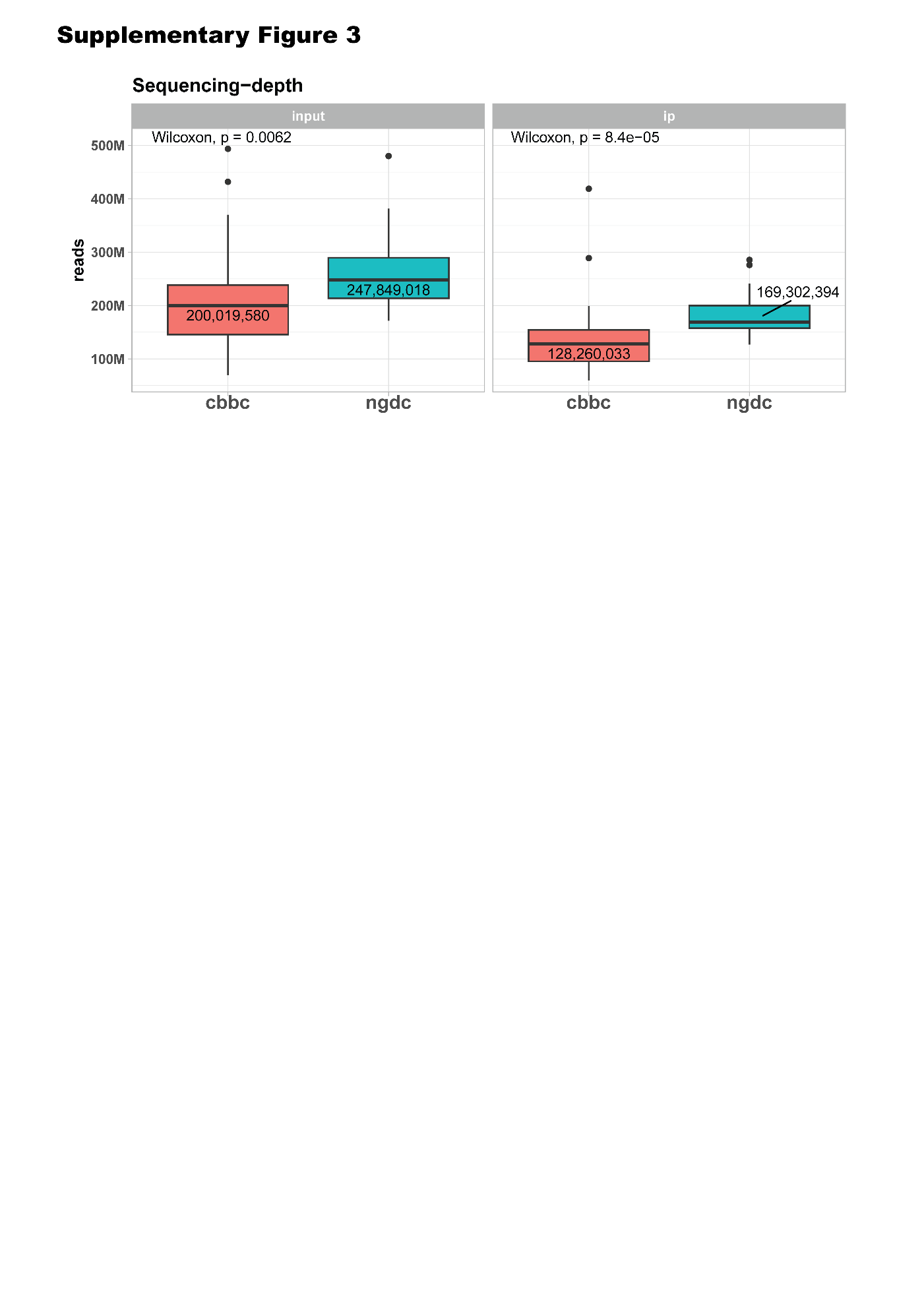

Supplementary Fig. 3 | Average number of sequence reads of input and ip sample in each dataset respectively.

In general, the sequencing-depth was significantly differed between the two datasets more in ip samples than input samples.

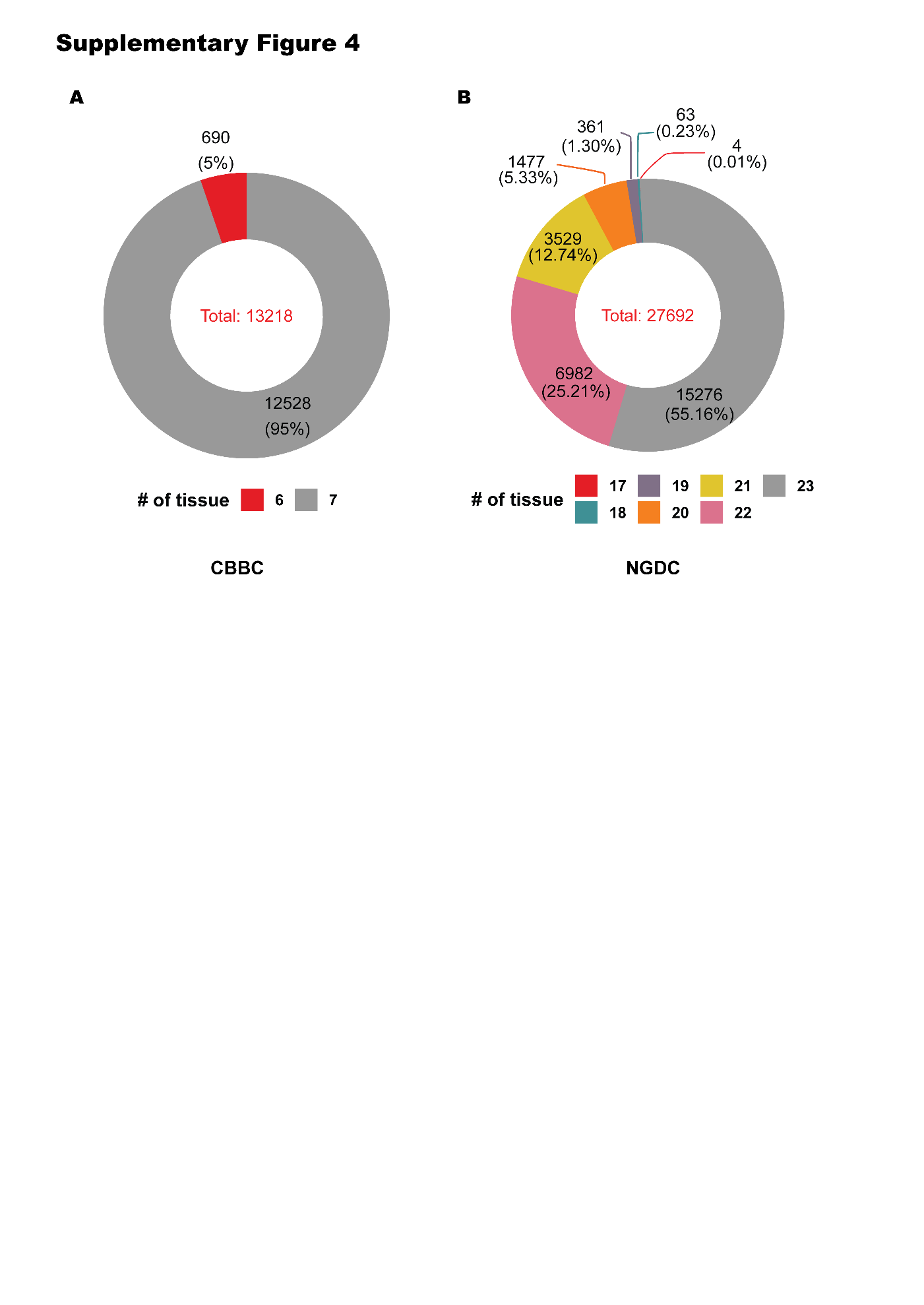

Supplementary Fig. 4 | The total number of significantly shared m6A sites (FDR<0.01) and the number of tissues that each site shared in each dataset.

(A) CBBC dataset. A total of 13,218 sites were significantly shared across tissue samples and 95% of them were shared in all 7 tissues in CBBC data. (B) NGDC dataset. A total of 27,692 sites were significantly shared across tissue samples and 93% of them were shared in nearly all 23 tissues (21~23) in NGDC data.

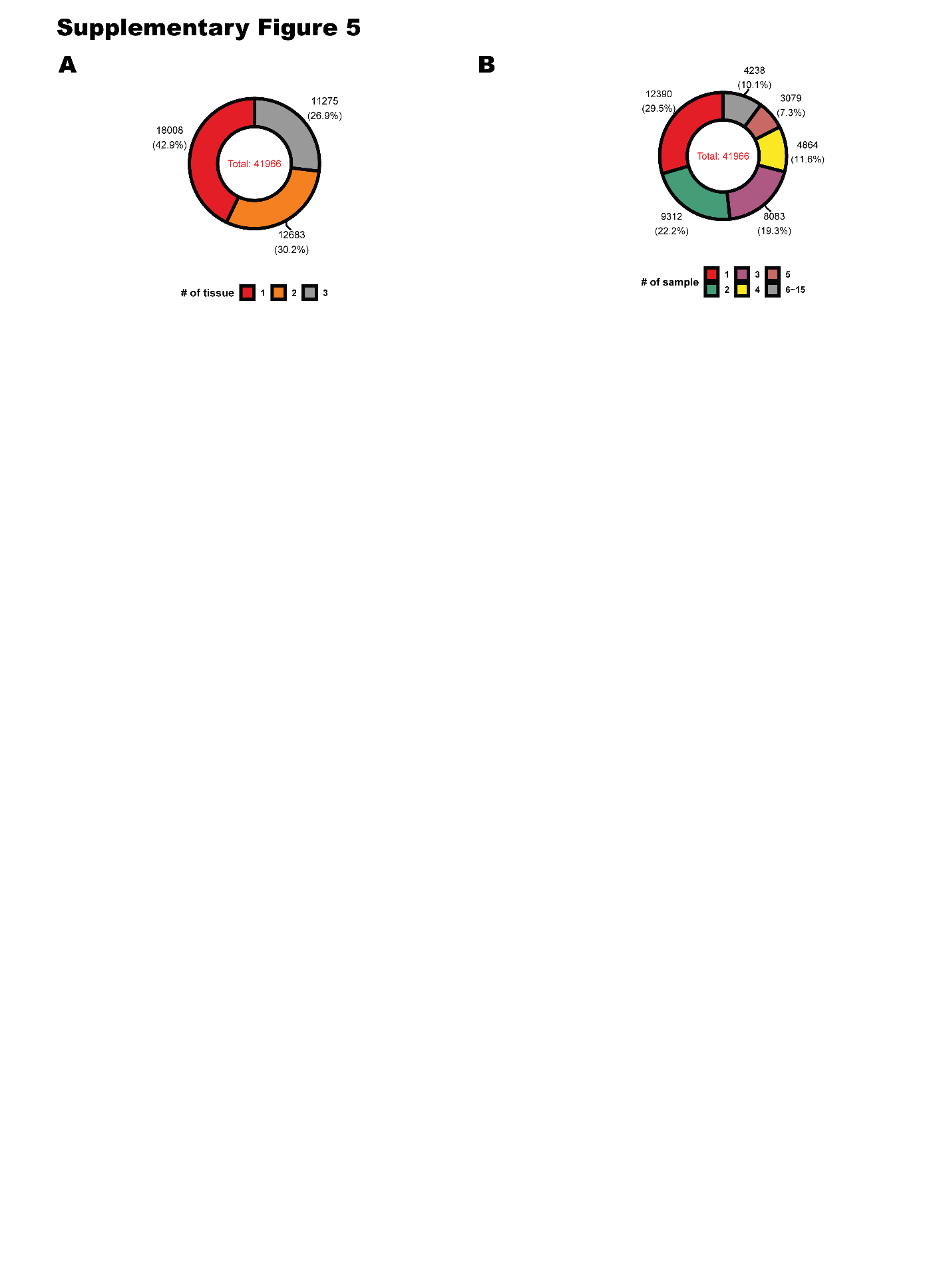

Supplementary Fig. 5 | Infrequent m6A sites. A total of 41,966 infrequent m6A sites that were methylated less than or equal to 3 tissues only.

(A) The number of tissues among infrequent sites. (B) The number of overlapping samples among infrequent sites. Most of the infrequent m6A sites (82.6%) were only presented in fewer than five samples.

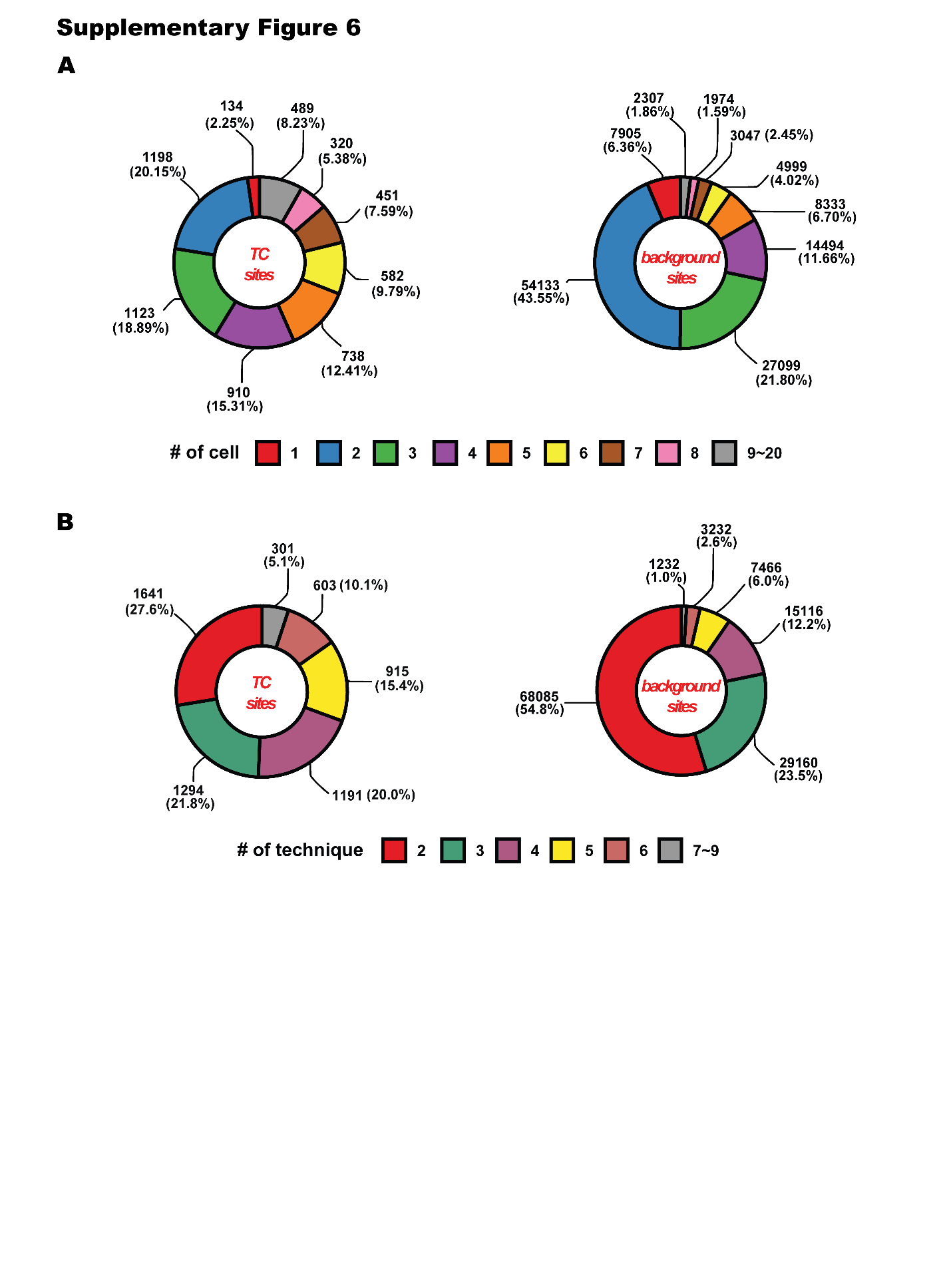

Supplementary Fig. 6 | Reproducibility and context independence of TC m6A sites and background m6A sites.

(A) The number of cell lines in which each site was identified. Approximately 58.7% of TC m6A sites were identified in ≥3 cell lines, compared with 28.3% of background sites. (B) The number of single-base m6A profiling methods by which each site was detected. While 72.4% of TC m6A sites were detected by ≥ 3 profiling methods, only 45.8% of background sites were similarly detected.

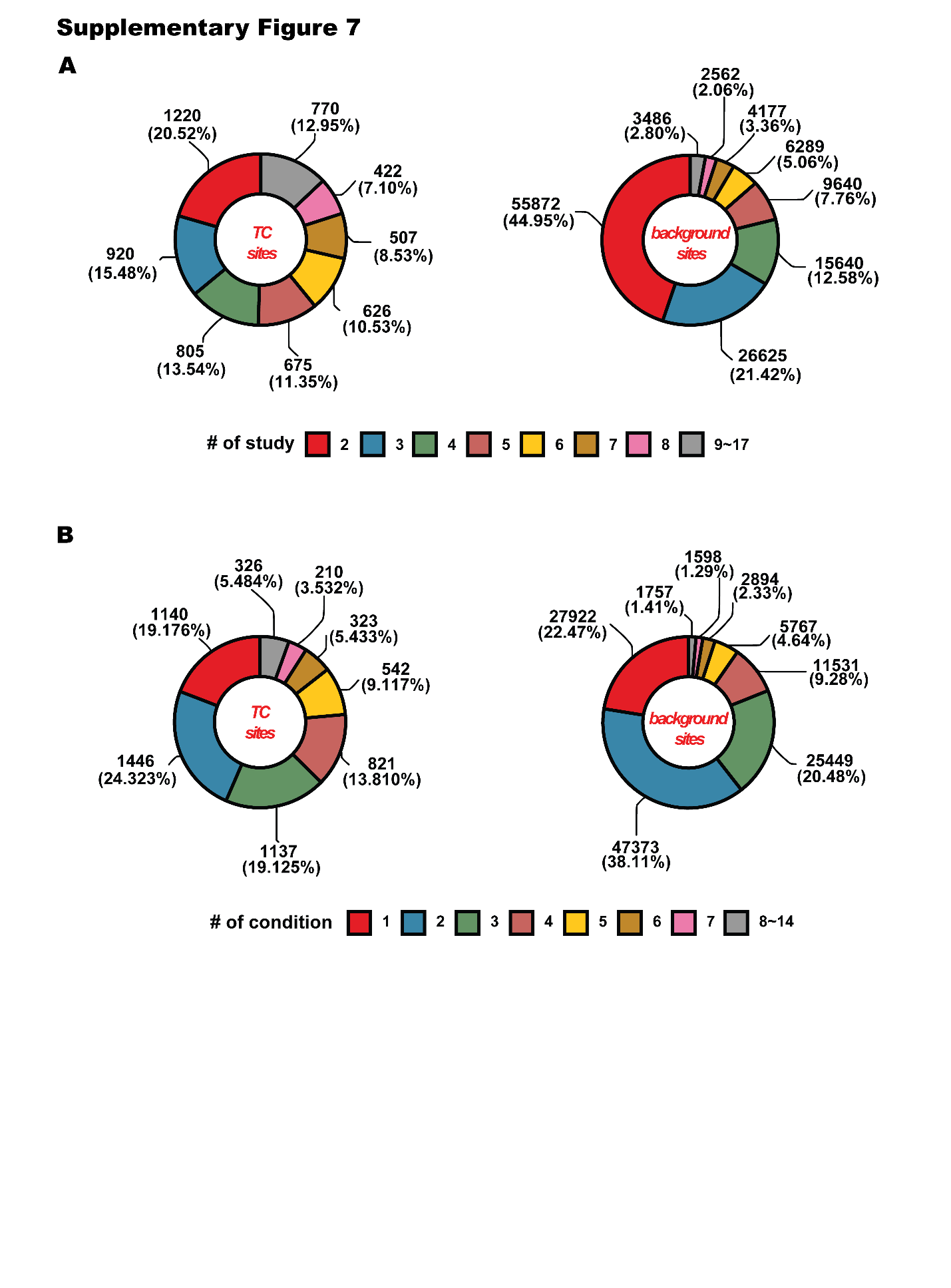

Supplementary Fig. 7 | Reproducibility and context independence of TC and background m6A sites.

(A) The number of independent studies reporting each m6A site. Approximately 50.5% of TC m6A sites were reported in more than four independent studies, compared with 20.1% of background sites. (B) The number of perturbed (non-normal) experimental conditions in which m6A sites were identified. Among TC m6A sites, 37.8% were identified in more than three perturbed conditions, whereas only 18.9% of background sites were identified in as many conditions.

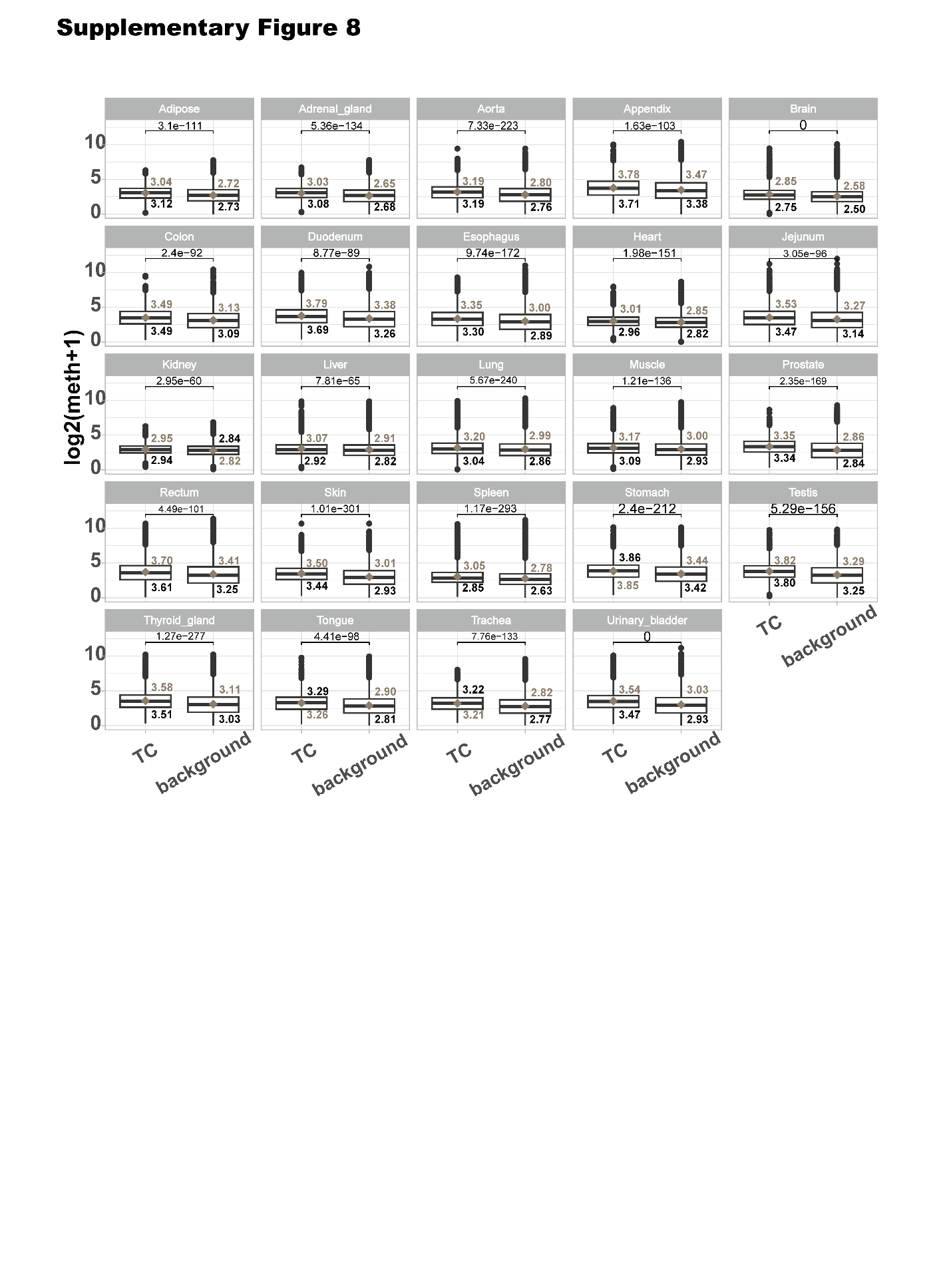

Supplementary Fig. 8 | Comparison of methylation levels between TC m6A sites and background m6A sites for each tissue respectively.

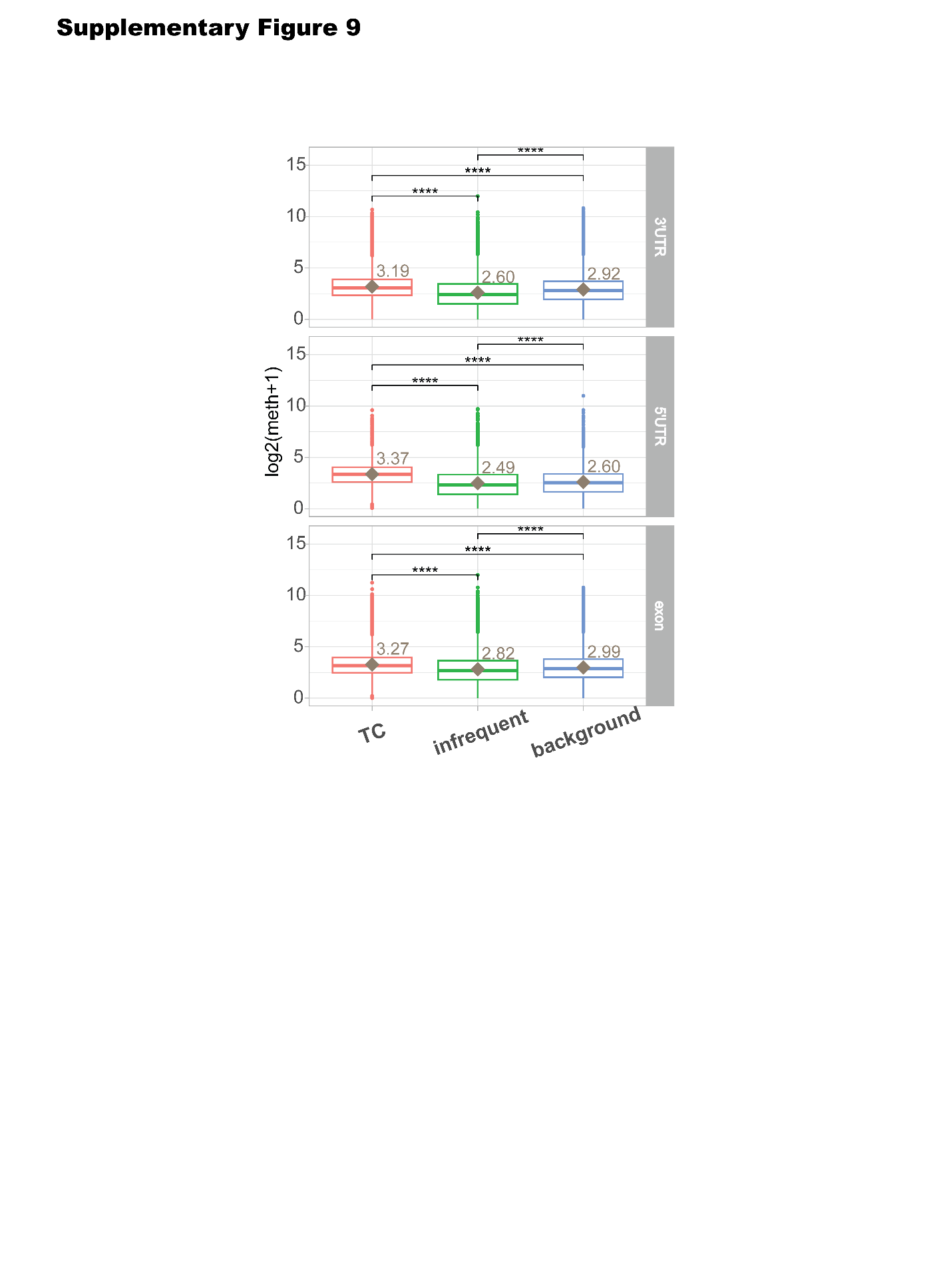

Supplementary Fig. 9 | Comparison of methylation levels between TC, infrequent, and background m6A sites in each exon, 3’UTR, and 5’UTR region respectively.

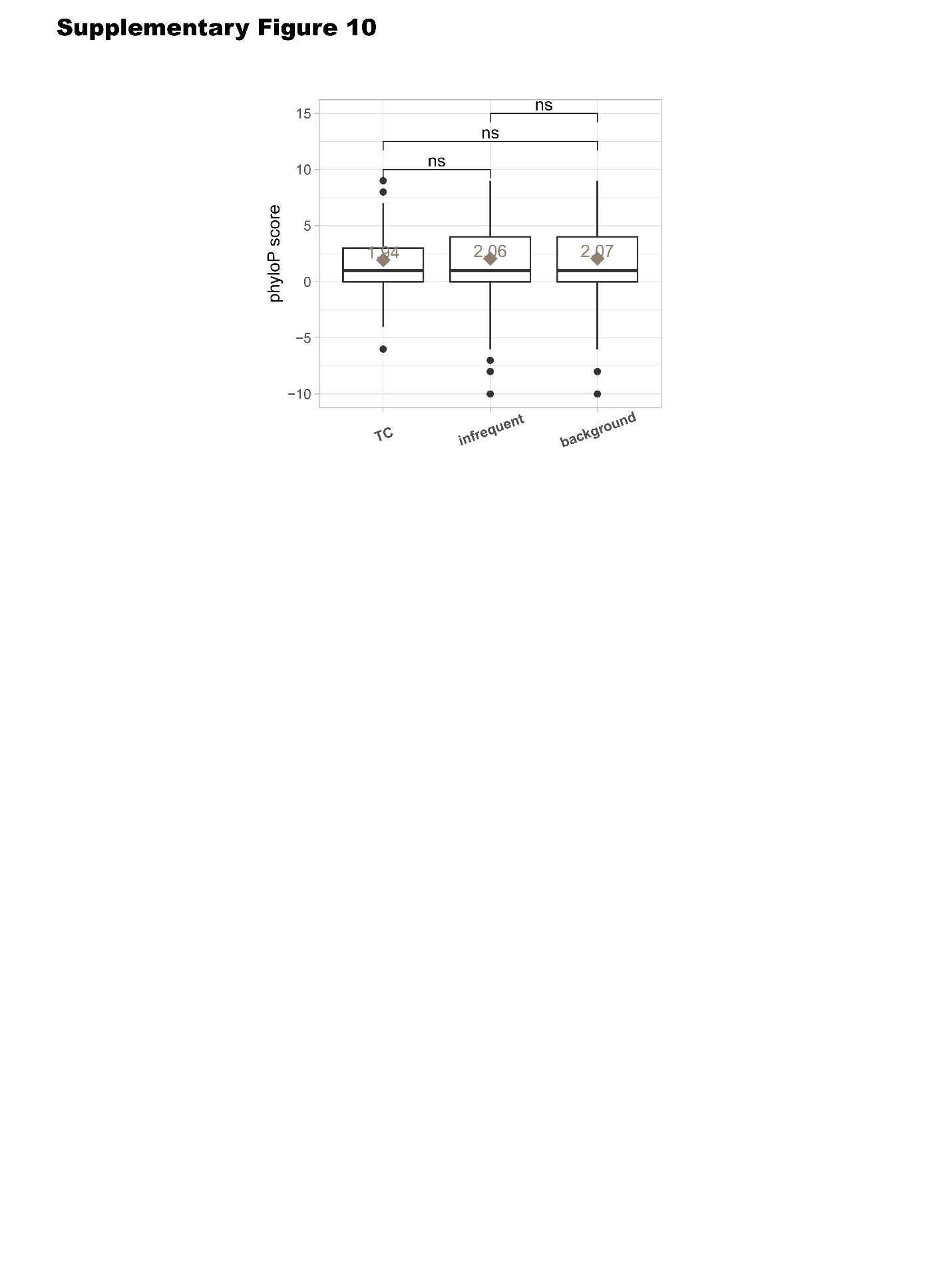

Supplementary Fig. 10 | Distribution of PhyloP conservation scores, demonstrating that m6A sites are localized in conserved or neutrally evolving regions.

TC, infrequent, and background m6A sites did not show statistical difference in PhyloP score, indicating that this localization is general feature of m6A deposition.

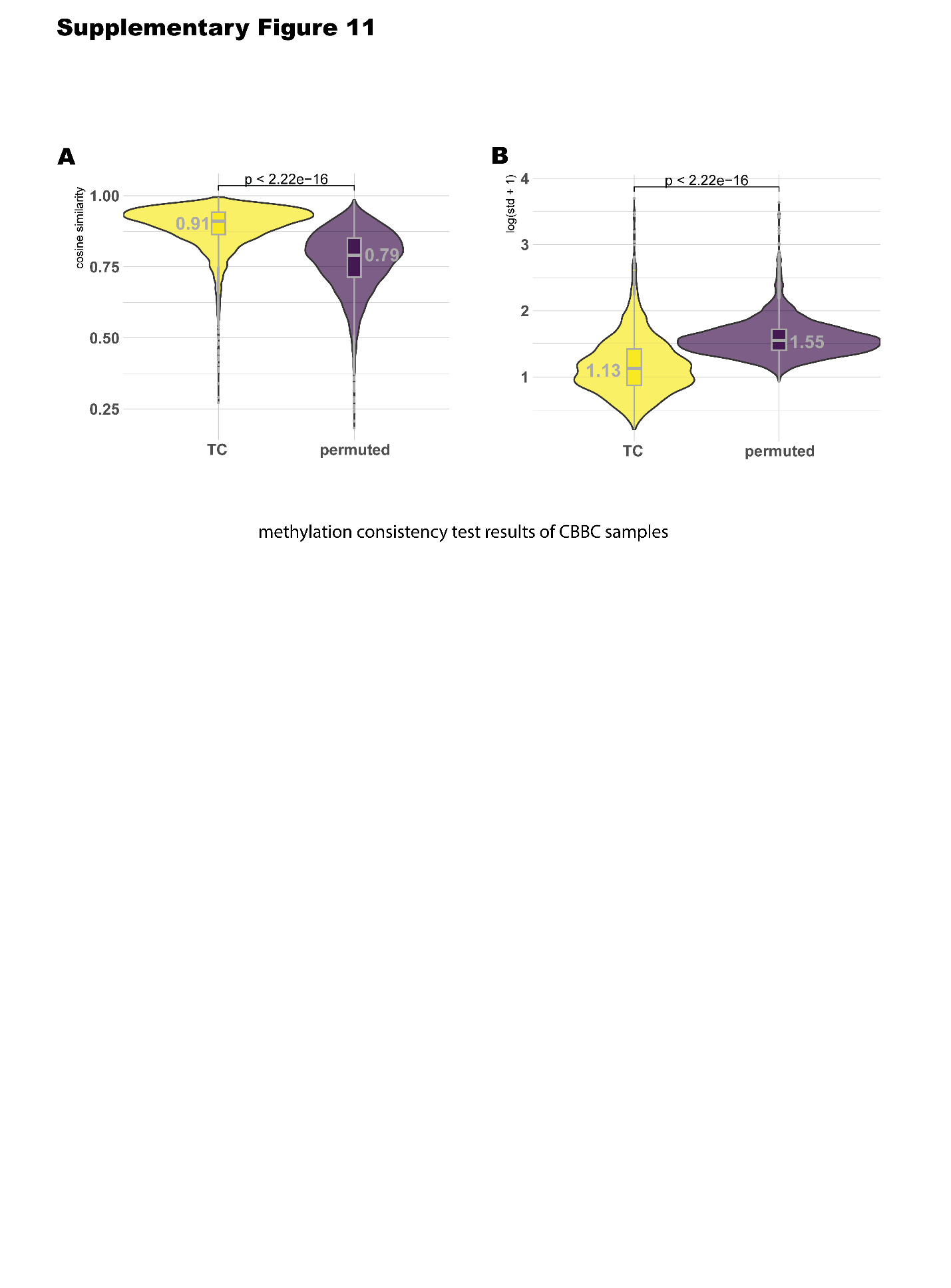

Supplementary Fig. 11 | Methylation consistency across tissues in CBBC samples.

(A) cosine similarity of methylation at TC sites across CBBC tissue samples exhibiting significantly higher similarity than expected under a random model (B) standard deviation of methylation at TC sites across CBBC tissue samples exhibiting significantly lower standard deviation in their methylation levels than expected under a random model.

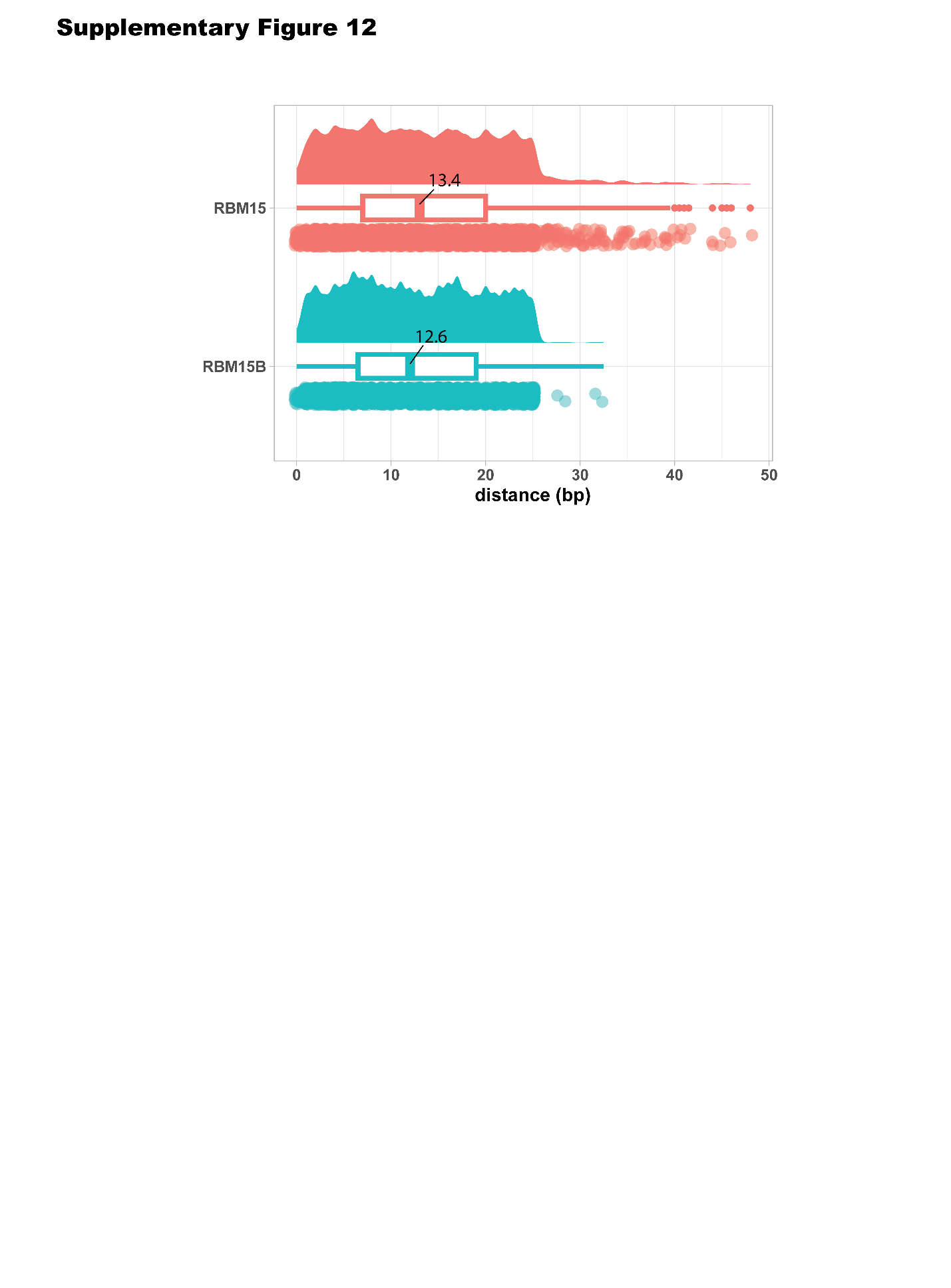

Supplementary Fig. 12 | The distribution of the center of binding peak regions.

RBM15 and RBM15B binding centers were located 13.4 bp (RBM15) and 12.6 bp (RBM15B) away from the TC sites on average.

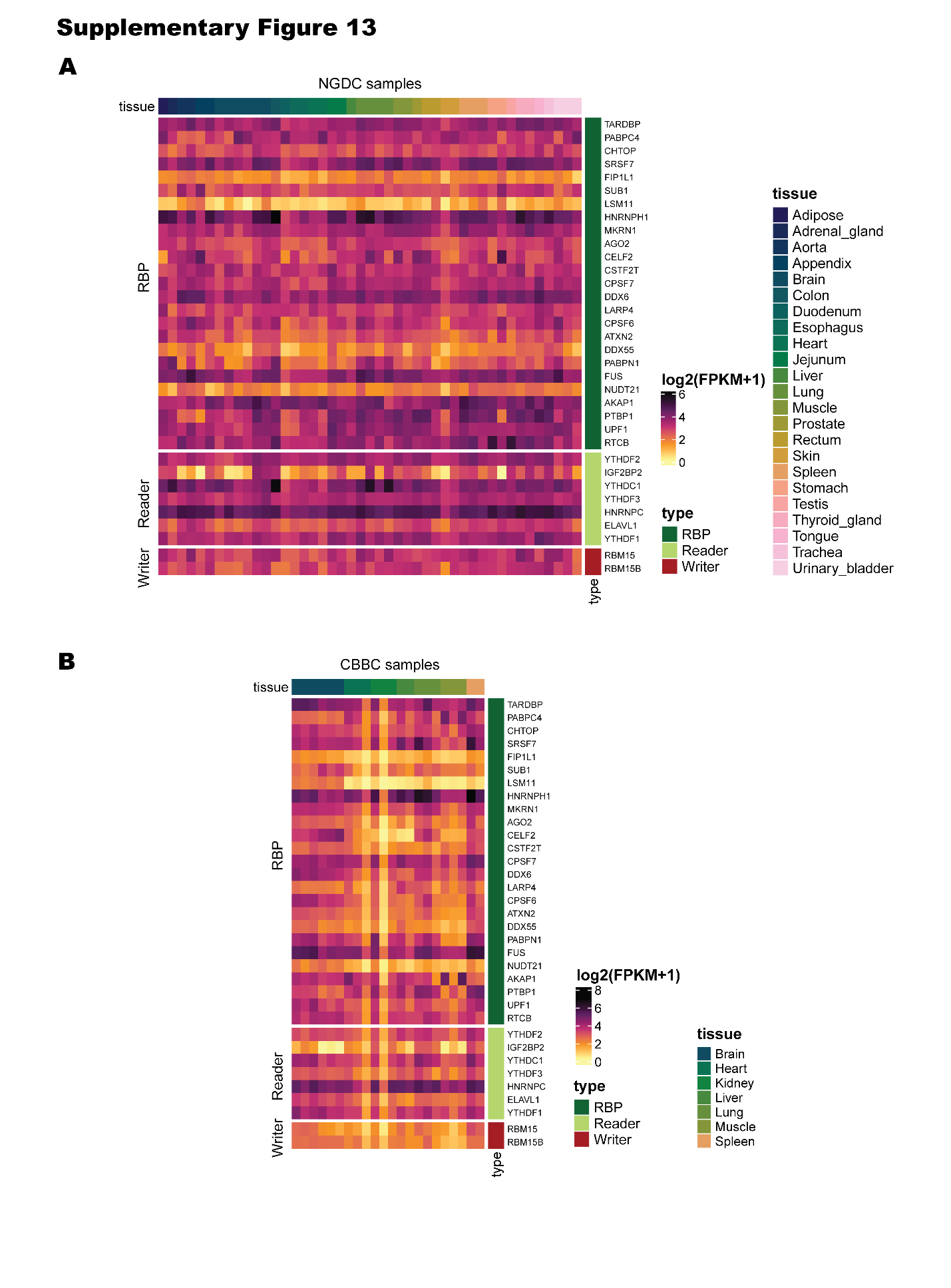

Supplementary Fig. 13 | Expression levels of RBPs across tissues in NGDC and CBBC data respectively.

Expression analysis revealed that both RBM15 and RBM15B are stably expressed across all tissues.

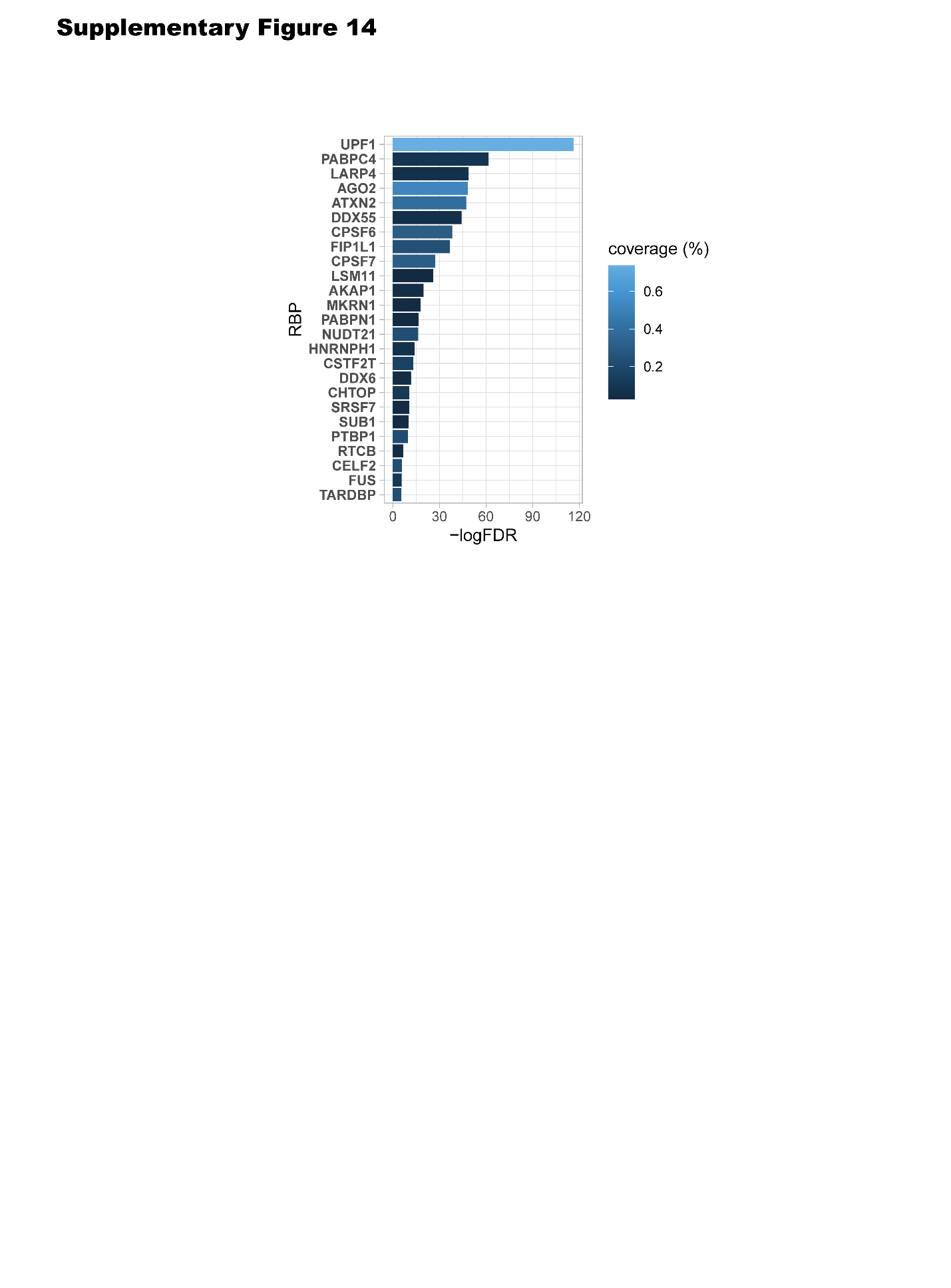

Supplementary Fig. 14 | 25 additional RBPs that are not m6A regulators (writer, eraser, or reader) but were also enriched near TC m6A sites

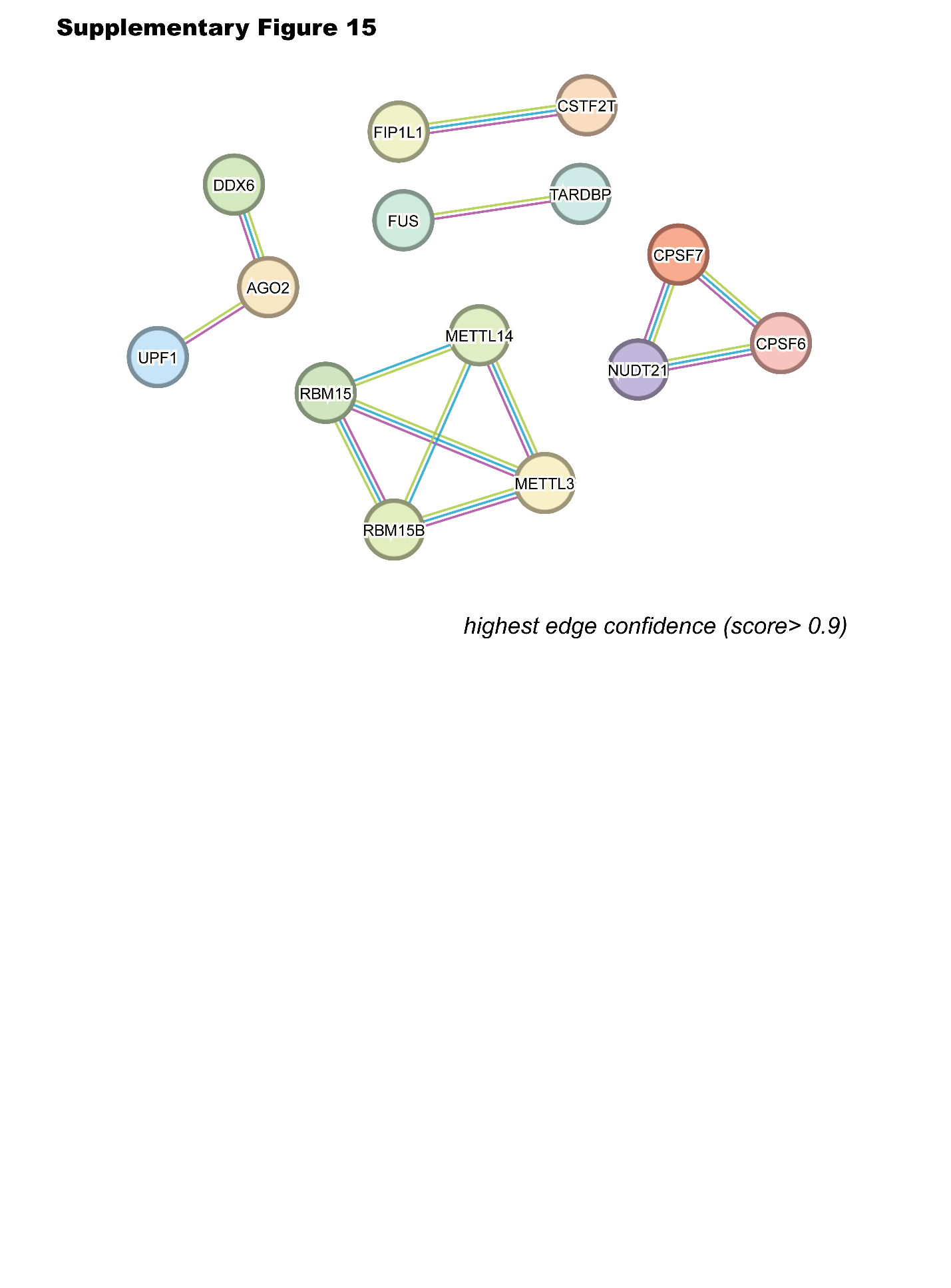

Supplementary Fig. 15 | protein–protein interaction analysis using STRING database.

This showed that only RBM15 and RBM15B directly interact with METTL3 or METTL14 (confidence score > 0.9).

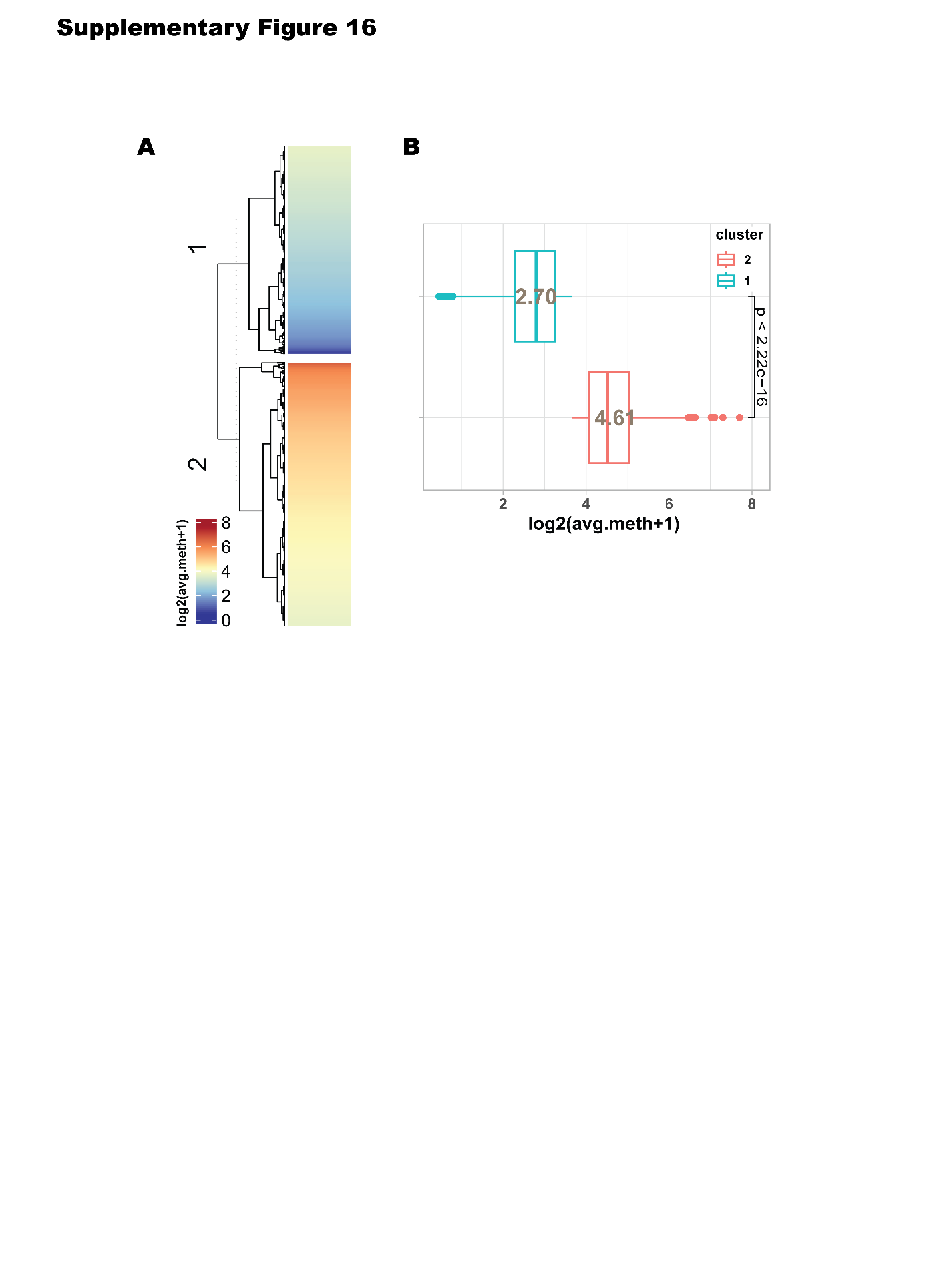

Supplementary Fig. 16 | Average methylation levels between high-methylation and low-methylation clusters from TC-m6A sites.

(A) Heatmap with clustergram of average methylation of TC site across tissue samples. (B) Statistical difference of average methylation between two clusters (high-meth cluster vs low-meth cluster)

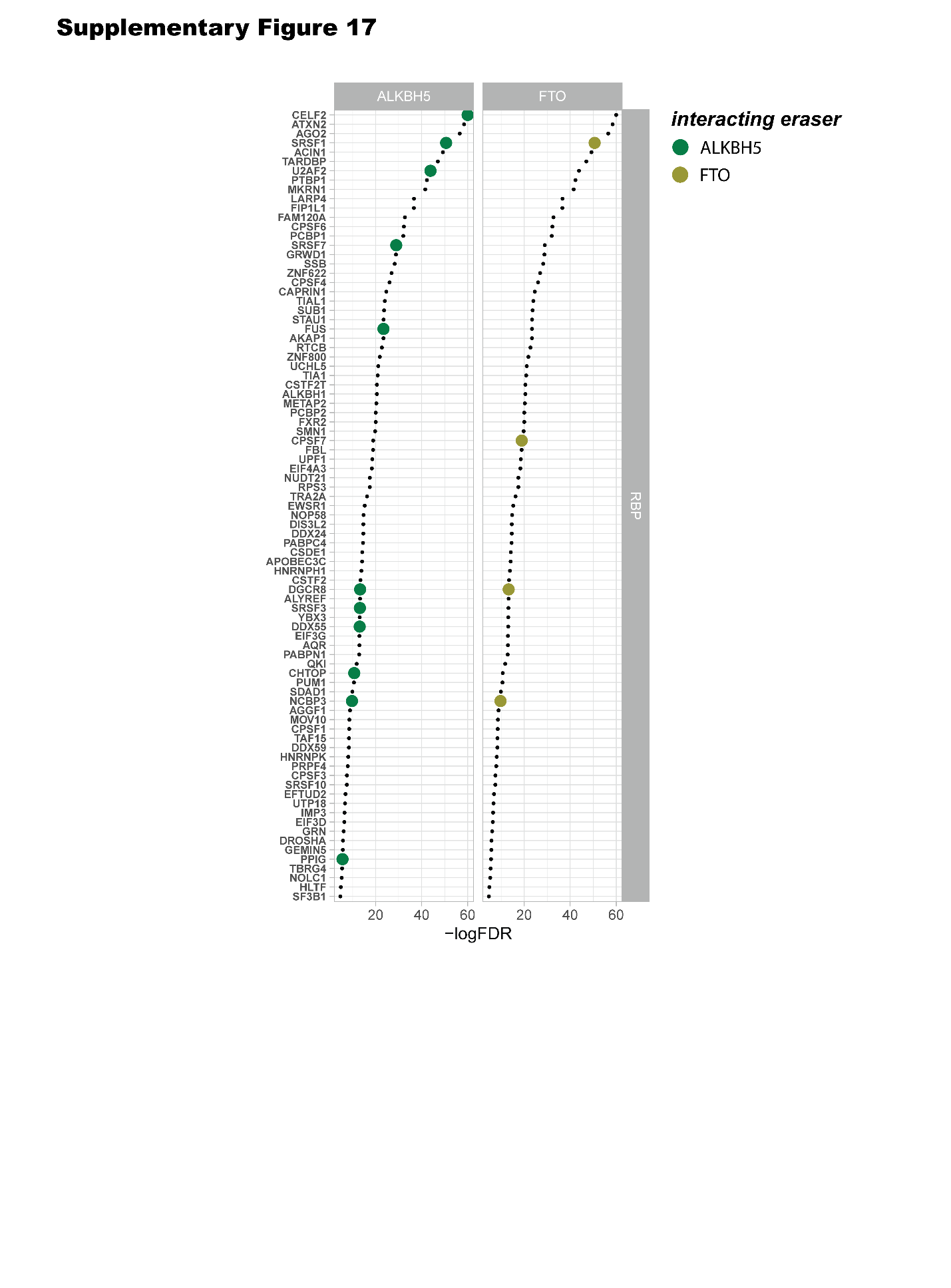

Supplementary Fig. 17 | 85 RBPs showed significant enrichment in the low-methylation cluster (Fisher’s exact test, FDR < 0.01).

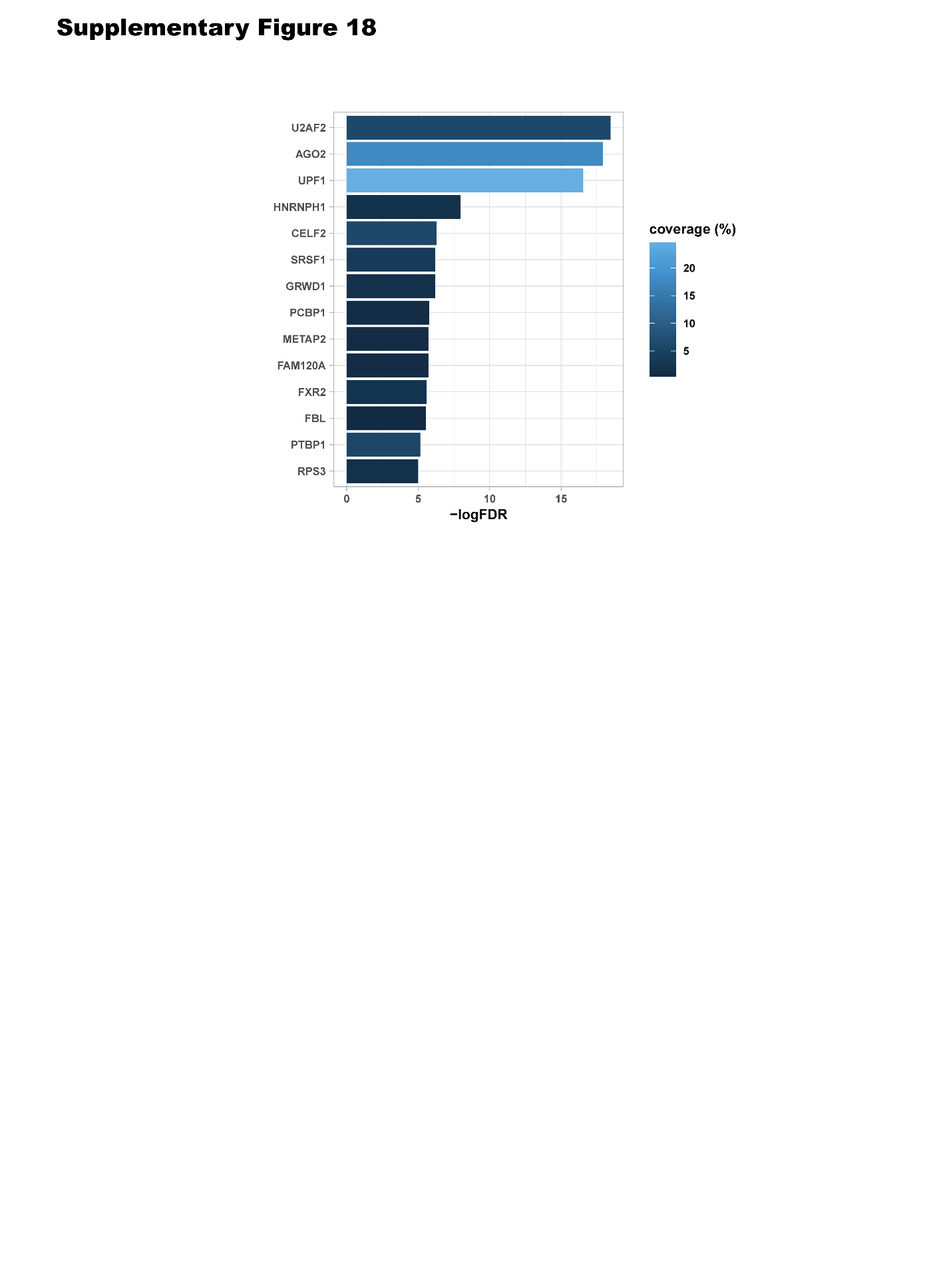

Supplementary Fig. 18 | Enriched RBPs observed in TC sites of the low-methylation cluster for ROI2

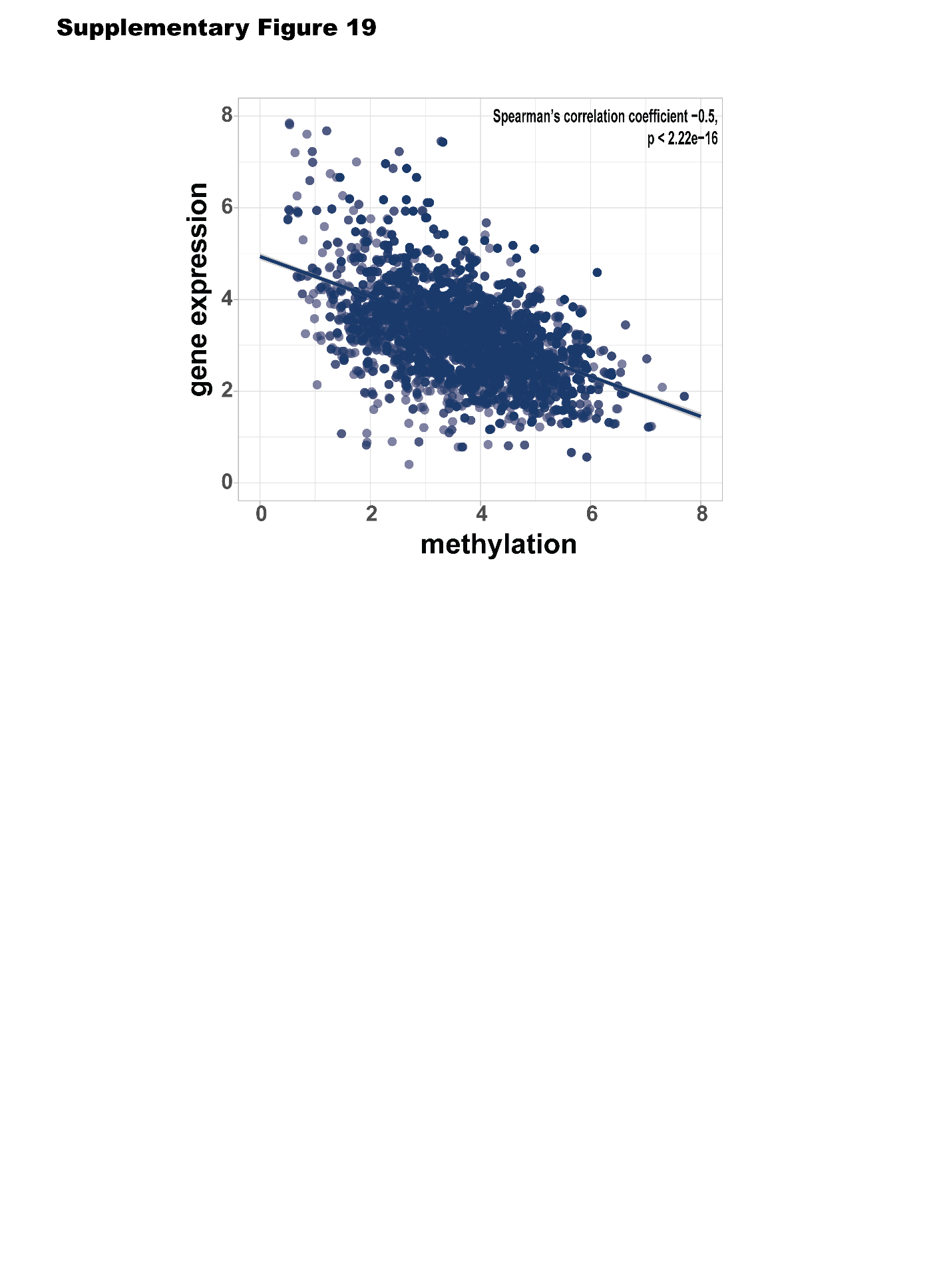

Supplementary Fig. 19 | Methylation levels of TC sites and expression levels of associated genes containing TC m6A sites in CBBC tissue samples.

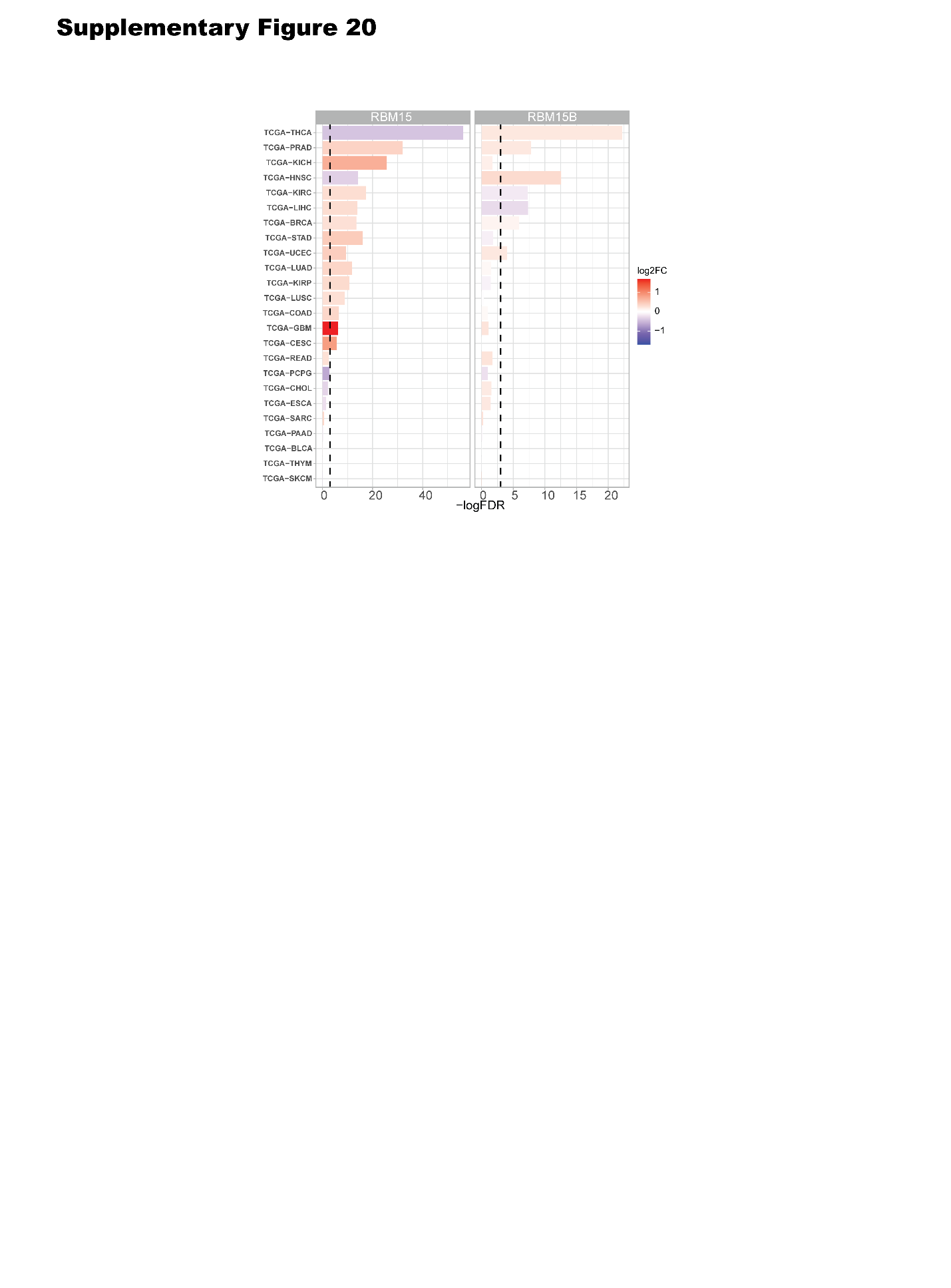

Supplementary Fig. 20 | Differential expression of the core TC m6A regulators RBM15 and RBM15B across TCGA cohorts. Dashed line represents FDR=0.05.
